# Higher rostral locus coeruleus integrity is associated with better memory performance in older adults

**DOI:** 10.1101/332098

**Authors:** Martin J. Dahl, Mara Mather, Sandra Düzel, Nils C. Bodammer, Ulman Lindenberger, Simone Kühn, Markus Werkle-Bergner

**Affiliations:** Center for Lifespan Psychology, Max Planck Institute for Human Development, Berlin, Germany; Davis School of Gerontology, University of Southern California, Los Angeles, USA; Max Planck UCL Centre for Computational Psychiatry and Ageing Research, London, England, and Berlin, Germany; Department of Psychiatry and Psychotherapy, University Clinic Hamburg-Eppendorf, Hamburg, Germany

**Author notes:** Correspondence concerning this manuscript should be addressed to MJD or MWB.

## Abstract

For decades, research into memory decline in human cognitive aging has focused on neocortical regions, the hippocampus, and dopaminergic neuromodulation. Recent findings indicate that the locus coeruleus (LC) and noradrenergic neuromodulation may also play an important role in shaping memory development in later life. However, technical challenges in quantifying LC integrity have hindered the study of LC-cognition associations in humans. Using high-resolution neuromelanin-sensitive magnetic resonance imaging, we found that individual differences in learning and memory were positively associated with LC integrity across a variety of memory tasks in younger (n = 66), and older adults (n = 228). Moreover, we observed functionally relevant age differences confined to rostral LC. Older adults with a more youth-like rostral LC also showed higher memory performance. These findings link non-invasive, in vivo indices of LC integrity to memory in aging and highlight the role of the LC norepinephrine system in the decline of cognition.

## Introduction

Memory performance declines with advancing adult age^1^, jeopardizing the everyday competence of older adults^2, 3^. Age-associated changes in neocortical regions, the hippocampus, and dopaminergic neuromodulation have been found to contribute to age-related memory impairments^1, 4–6^. More recently, findings from animal and post-mortem human research have led researchers to propose that cell loss and accumulation of abnormal tau in the locus coeruleus (LC), the brain’s primary norepinephrine (NE) source, are critically related to cognitive decline in normal aging and age-related pathologies^7–10^. However, direct in vivo evidence relating LC integrity to age differences in general memory abilities in humans is scarce^11^ (for evidence from animal models, see for example^12–14)^. While there is an initial indication of a selective role of LC integrity, as indexed by magnetic resonance imaging (MRI; see below), in the encoding of negative emotional events^15^, the relation of LC integrity to memory performance in general is still an open question. Given LC’s pivotal role in age-related memory disorders like Alzheimer’s disease^8, 9, 16^, this relationship, however, is of high clinical and scientific importance.

The LC is a hyperpigmented cylindrical cluster of catecholaminergic neurons located in the dorsorostral tegmentum^17^. Roughly symmetrical in both hemispheres, it extends only about 15 mm rostrocaudally from the level of the inferior colliculi to a position in the lateral wall of the fourth ventricle^18, 19^. Despite its small size, the LC has diffuse and highly arborized efferent projections throughout the brain^20^. While initial reports assumed the LC was a uniform structure, current evidence indicates a spatially differentiated organization. Cells giving rise to dense hippocampal projections tend to be located in more rostral segments while those that innervate the cerebellum and spinal cord are located more caudally^21–23^. NE release by the LC modulates cognitive functions such as perception, attention, learning and memory^21, 24–32^. In particular, via its action on ß-adrenoceptors in the hippocampus, the LC modulates long-term potentiation (LTP) and long-term depression (LTD), key determinants of synaptic plasticity and memory^33, 34^. Recent optogenetic work has confirmed a causal role of LC activity in hippocampal LTP and memory enhancement, potentially via co-release of dopamine^35–37^.

Its location adjacent to the ventricular system and its widespread, unmyelinated projections expose LC cells to blood- and cerebrospinal-fluid bound toxins, making them a likely target for neurodegeneration^11^. Consistent with its age-related vulnerability, a prominent, rostrally accentuated decline of LC density was reported in healthy aging (see Manaye and colleagues^22^ for a review of early studies; but see^16, 18, 38, 39^ and Mather and Harley^11^ for a recent discussion). Based on longitudinal data, Wilson and colleagues^10^ reported initial evidence pointing to the importance of LC integrity, here assessed via LC’s neuronal density, for maintaining memory abilities in old age. The authors evaluated cognitive abilities in a sample of older adults annually over a mean duration of about six years and, upon the participants’ death, assessed neuronal density in the LC via autopsy. Wilson and colleagues^10^ observed an attenuation of cognitive decline as well as higher baseline cognitive abilities in individuals with higher LC integrity even after accounting for the integrity of other neuromodulatory systems and markers of neuropathology (i.e., Lewy bodies and neurofibrillary tangles within the brainstem)^10^. Furthermore, studies in aged mice^12^, rats^13^, and monkeys^14^ in which NE or its agonists were manipulated indicate that NE plays a causal role in learning and memory.

Non-invasive, in vivo assessments of human LC integrity are notoriously difficult, given the nucleus’ small size and location deep in the brainstem^30, 40, 41^. Fortunately, however, a by-product of catecholamine synthesis, the dark, insoluble pigment neuromelanin, accumulates in the LC across the life span^42^. LC pigmentation increases from birth until late middle age^42^. While some studies observed constant neuromelanin levels in late life^38^, others reported a decline from midlife until death, probably due to selective atrophy of neuromelanin-containing cells^22, 43, 44^. In noradrenergic cells, neuromelanin avidly chelates metals such as copper and iron and, as a compound, shows paramagnetic, T1-shortening effects^45^. Sasaki and colleagues developed a neuromelanin-sensitive Turbo Spin Echo (TSE) MRI sequence that visualizes the LC as an hyperintense area adjacent to the lateral floor of the fourth ventricle^46^ (for a review of neuromelanin-sensitive MRI studies, see Liu and others^47^). Keren and colleagues^48^ recently validated this sequence by scanning human post-mortem samples using ultra high field MRI and then performing histological analyses of the samples. Brainstem areas showing hyperintensities on neuromelanin-sensitive scans overlapped closely with noradrenergic cells as identified by histology. Thus, neuromelanin acts as a natural contrast agent that opens the door to the non-invasive, in vivo assessment of LC integrity via MRI.

To determine the importance of MRI-indexed, structural LC integrity for the maintenance of memory functioning in vivo, we assessed individual differences in learning and memory among younger and older adults with the Rey Auditory Verbal Learning Test (RAVLT), a validated neuropsychological measure of memory functioning. In this context, specifically the analysis of individual learning trajectories, that is, the increase in recall performance across iterative item presentations, conveys valuable information about a participant’s current and future cognitive status^49–52^. Using structural equation modeling (SEM) to capture the non-linear dynamics of growth in performance^53^, previous studies have linked aging to lower initial memory performance^54, 55^ and slower learning with practice^55^. Hence, we hypothesized that the integrity of the LC-NE system as assessed by neuromelanin-sensitive MRI would be closely associated with individual differences in initial memory performance and learning rates. In sum, the goal of this study was to extend our knowledge about the role of the LC-NE system in human cognitive aging by linking non-invasive, in vivo indices of LC integrity to memory abilities in younger and older adults.

## Results

### Initial recall performance is lower in older adults

We assumed that younger and older adults recall a gradually increasing number of words over iterative RAVLT learning trials (see Figure 1 and Methods, section on cognitive data assessment). Formally, this can be expressed as a learning curve consisting of an initial memory performance level (intercept) and a gain over learning trials (slope). Since we hypothesized that the participants’ learning performance reaches a natural (or task-induced) performance limit, we expected an increasing yet negatively accelerated slope^53, 54^. Using latent growth curve modeling (LGCM, a specific variant of SEM), we estimated the intercept and slope factors on a latent level^56^. To test for differences in intercept and slope parameters between age groups, we opted for a multiple-group model^57, 58^. In particular, we fitted a simple quadratic growth model comprising an intercept as well as quadratic and linear slope factor for each age group (see Supplementary Figure 3). To investigate potential relations between intercept and slope terms (e.g., starting out higher may be related to faster performance increases), we freely estimated covariances between intercept and slope parameters within age groups^55^.

**Figure 1.**
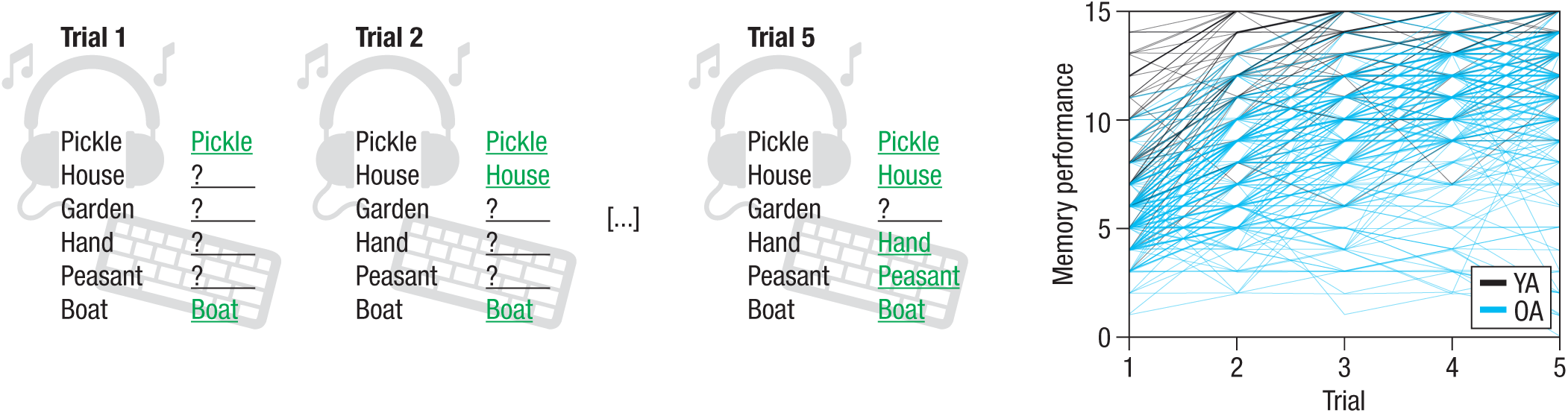
Schematic overview of the verbal learning and memory task (left). A word list consisting of 15 unrelated words is auditorily presented to participants over five trials. After each trial, subjects freely recall items and enter their responses via keyboard. The sum of correctly recalled words provides the performance measure for each trial (ranging from 0–15). Right: Younger (YA; n = 66; black) and older adults’ (OA; n = 228; blue) performance increases over learning trials in a non-linear fashion.

The adequacy of the proposed model was assessed using two frequently reported fit indices: First, the root mean square error of approximation (RMSEA) is a closeness of fit coefficient that expresses how much the postulated model approaches the true model. Second, the comparative fit index (CFI) is an incremental fit index which compares the goodness of fit of the proposed model with a more restrictive nested baseline model^57, 59, 60^. RMSEA values close to or below 0.06 and CFI values of close to 0.95 or greater indicate good model fit^59^.

A multiple-group model describing the gradually increasing but negatively accelerated performance trajectories as a quadratic growth function fit the data well and outperformed competing alternative models (*χ*^2^(46) = 81.764, RMSEA = 0.052, CFI = 0.965^59^; see Supplementary Results 2.1.1, and lower part of Figure 2). Importantly, the model allows testing of group differences in factors capturing initial recall (intercept) and learning (slope) as well as for interindividual differences therein within each age group. Parameters of interest were evaluated using likelihood ratio tests. In particular, constraining parameters to be equal (to test between-group differences) or to zero (for within-group comparisons) should result in significant decrease in model fit relative to a model with unconstrained parameters in case of reliable differences^58^. The difference in model fit between the constrained and the unconstrained model (i.e., Δ*χ*^2^) under the null hypothesis follows a *χ*^2^-distribution with the degrees of freedom equivalent to the difference in numbers of constrained parameters (i.e., Δ*df*)^58^.

**Figure 2.**
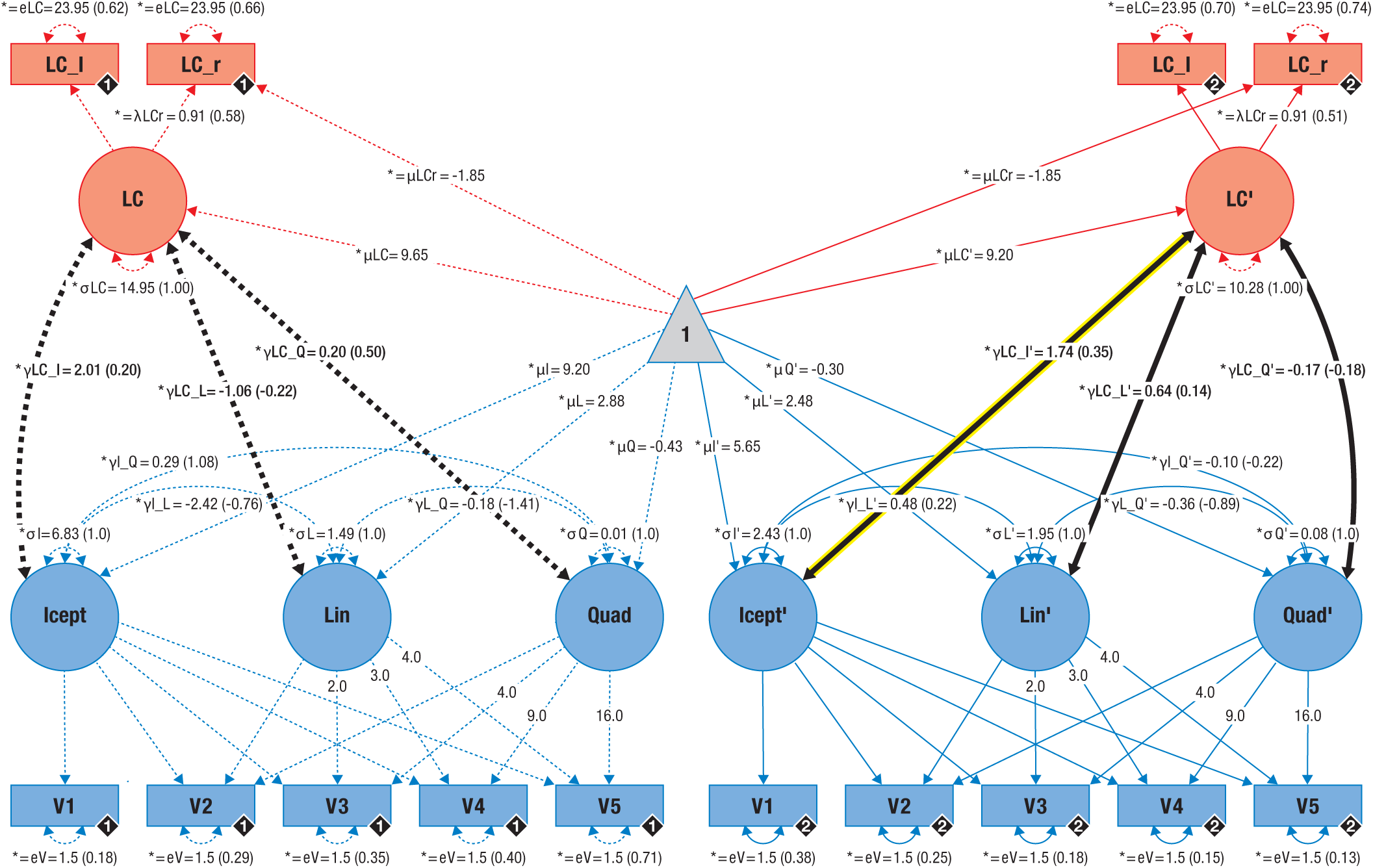
Pictorial rendition of the structural equation model that probes associations (thick black lines) between locus coeruleus integrity (LC; red) and memory performance (blue) in younger and older adults on a latent level. Cognitive manifest variables represent the iteratively assessed memory performance in a verbal learning and memory task (V1–5). Neural manifest variables are the LC intensity ratios of each hemisphere (left = LC_l; right = LC_r). Black diamonds on manifest variables indicate the age group (younger adults = 1, (n = 66), broken lines; older adults = 2, (n = 228), solid lines). (Co)Variances (*γ, σ*) and loadings (*λ*) in brackets indicate standardized estimates. Loadings that are freely estimated (*) but constrained to be equal across age groups (=) are indicated by both asterisk and equal signs (*=). Icept = Intercept; Lin/Quad= linear / quadratic slope, respectively. Rectangles and circles indicate manifest and latent variables, respectively. The constant is depicted by a triangle. The significant association between locus coeruleus integrity and memory performance (Icept) in older adults is highlighted (yellow frame).

Both the means and the variances of the intercept and slope factors differed reliably from zero in each age group (likelihood ratio tests for means: all Δ*χ*^2^(Δ*df* = 1) ≥ 62.602, all *p* < 0.001; likelihood ratio tests for variances: all Δ*χ*^2^(Δ*df* = 1) ≥ 11.97, all *p* < 0.001; see Supplementary Results 2.1.2). As the sole exception, there were no reliable interindividual differences in the variance of the quadratic slope factor in younger adults (likelihood ratio test: Δ*χ*^2^(Δ*df* = 1) = 0.276, *p* = 0.599, estimate (est) = 0.011, [95% confidence interval (CI): −0.031, 0.053]). Performance differences between younger and older adults were mainly found in intercepts (likelihood ratio test: Δ*χ*^2^(Δ*df* = 1) = 59.533, *p* < 0.001, est_younger adults_ = 9.197, [95% CI: 8.508, 9.886], est_older adults_ = 5.647, [95% CI: 5.395, 5.899], see Supplementary Results 2.1.3). In sum, learning rates did not differ reliably between age-groups, but older adults started and ended with lower recall scores compared to younger adults due to differences in initial recall.

### Locus coeruleus integrity scores are positively associated with initial recall performance in older adults

We developed a semi-automatic procedure to extract individual LC ratios^47^, that is, a ratio score of peak LC MRI intensity relative to peak intensity in a dorsal pontine reference region, across the rostrocaudal extent of the nucleus (see Figure 3a–c and Methods, section on magnetic resonance imaging data analysis). The procedure evinced high reproducibility across multiple measurements (see Supplementary Results 2.2.1.3) and was validated using both published LC maps (see Figure 3d) and manual intensity assessments (see Supplementary Results 2.2.1.2). In order to relate memory performance to LC integrity, we integrated LC ratios over slices and hemispheres to derive a single measure reflecting LC integrity. In particular, we estimated latent scores for LC integrity by means of a multiple-group single-factor structural equation model based on each hemisphere’s mean intensity ratios (see upper part of Figure 2 and Methods, section on estimation of latent locus coeruleus integrity scores). The proposed model fit the data well (*χ*^2^(7)= 1.791, RMSEA = 0.000, CFI = 1.222^59^; see Supplementary Results 2.2.3.1). We detected reliable average LC scores for both age groups; moreover, within each group, participants showed significant interindividual differences in LC scores (likelihood ratio tests: all Δ*χ*^2^(Δ*df* = 1) **≥** 18.305, all *p* **<** 0.001; see Supplementary Results 2.2.3.2). Younger and older adults showed no reliable differences in average LC scores (likelihood ratio test: Δ*χ*^2^(Δ*df* = 1) **=** 0.383, *p* = 0.536, est_younger adults_ = 9.640, [95% CI: 8.342, 10.939], est_older adults_ = 9.205, [95% CI: 8.484, 9.926]), in line with spatially confined age differences across the rostrocaudal LC axis^41, 61^.

**Figure 3.**
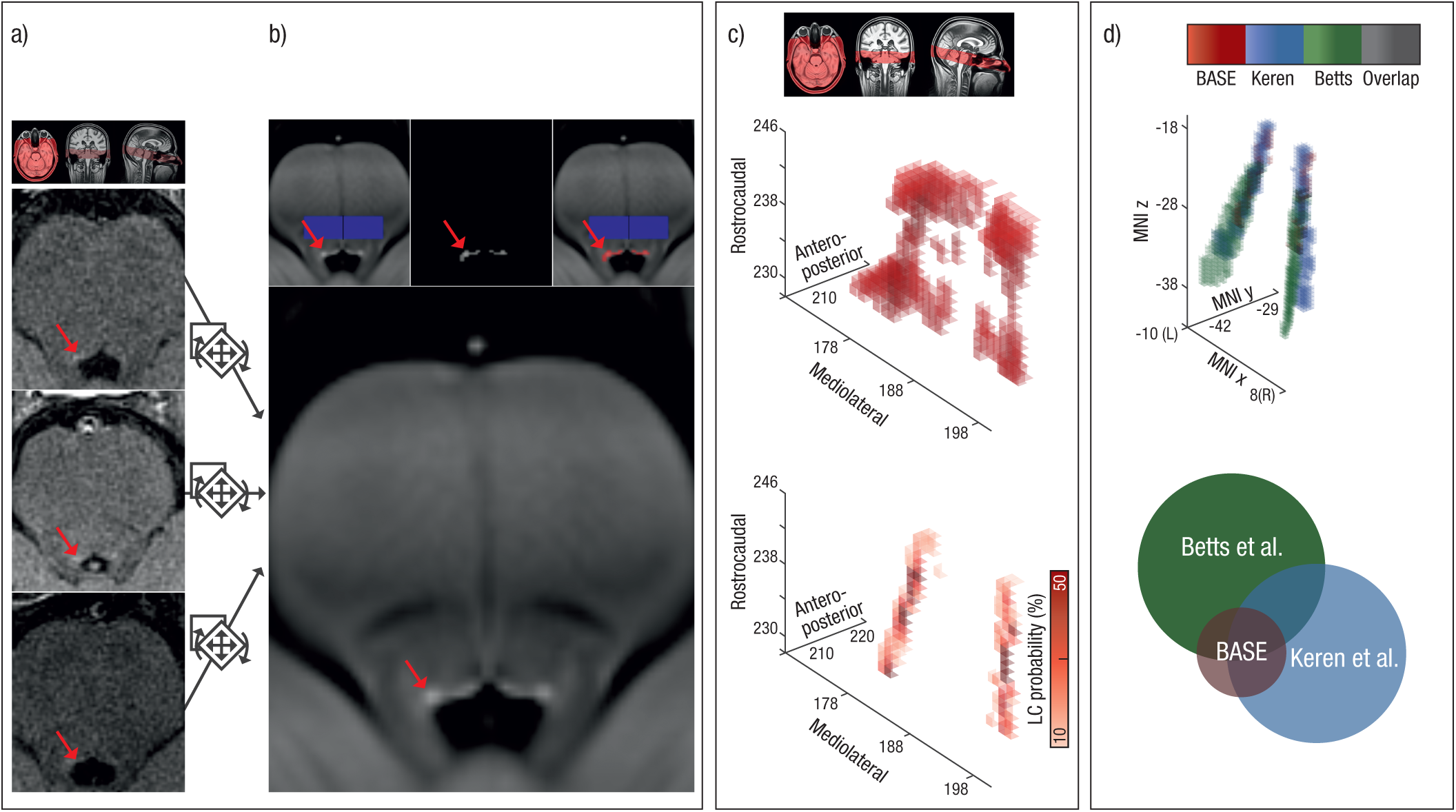
Schematic overview of the semi-automatic analysis procedure used to extract individual locus coeruleus (LC) intensity values across the rostrocaudal extent. (a) Native space neuromelanin-sensitive brainstem scans of three randomly selected subjects (axial slices are shown). Hyperintensities corresponding to the LC are indicated by red arrows. (b) Neuromelanin-sensitive scans were aligned and pooled across subjects in order to increase signal-to-noise ratio and facilitate LC delineation using a template-based approach. On a group level, LC location (red) was semi-automatically determined based on an intensity threshold relative to a pontine reference area (blue; see inlays). (c) Areas surviving the thresholding are grouped into a volume of interest (search space: upper plot; 3D representation) and used to restrict automatized extraction of individual peak intensities and their location. Observed peak LC locations were converted to a LC probability map (lower plot). (d) In standard space, the LC probability map was successfully validated using published maps by Keren and colleagues (2009)^41^ and Betts and colleagues (2017)^61^. Circle radius indicates map size (i.e., number of voxels). We freely share the resulting LC probability map here: https://www.mpib-berlin.mpg.de/LC-Map

After establishing valid models for both memory performance and LC integrity in isolation, we aimed to combine the independent information from both modalities. For this, we merged both SEMs in a unified neuro-cognitive model that demonstrated good fit (*χ*^2^(87) = 101.942, RMSEA = 0.024, CFI = 0.986^59^; see Figure 2 and Supplementary Results 2.3). Initial recall (intercept) was positively related to latent LC scores for older adults while the association failed to reach significance for the younger group (likelihood ratio test; for older adults: Δ*χ*^2^(Δ*df* = 1) **=** 7.939, *p* = 0.005, est_older adults_ = 1.737, [95% CI: 0.447, 3.027], standardized est_older adults_ = 0.348; for younger adults: Δ*χ*^2^(Δ*df* = 1) = 1.181, *p* **=** 0.147^15^, est_younger adults_ = 2.014, [95% CI: −1.671, 5.698], standardized est_younger adults_ = 0.199). Learning rates (slope) were not reliably associated with LC scores (likelihood ratio tests: all Δ*χ*^2^(Δ*df* = 1) **≤** 1.426, all *p* **≥** 0.232; see Supplementary Table 8). These findings indicate higher initial recall performance (intercept) for older adults with high LC integrity than for those with low LC integrity (see Figure 4).

**Figure 4.**
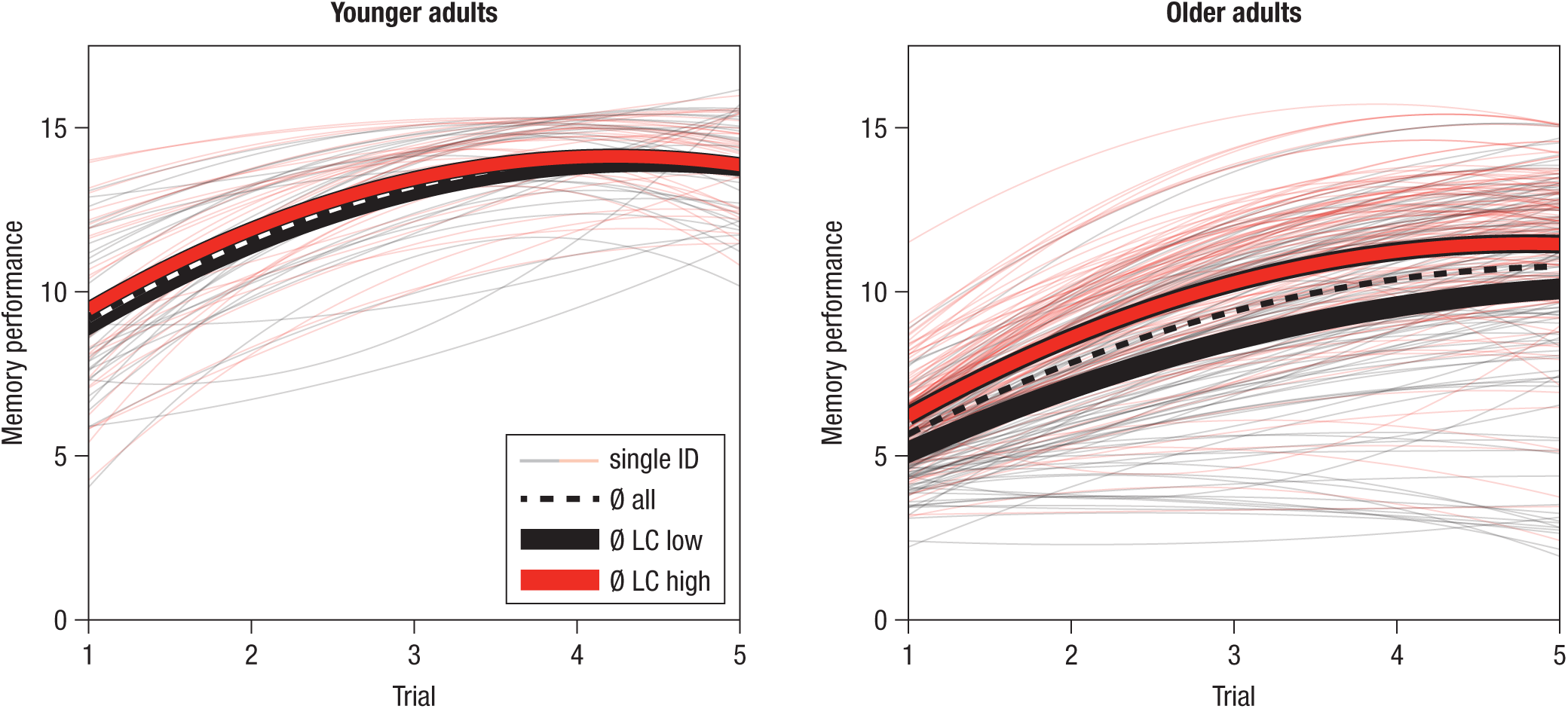
Estimated learning and memory performance trajectories for younger (left panel; n = 66) and older adults (right panel; n = 228). For visualization of the association between locus coeruleus (LC) integrity and memory performance, single subjects (thin lines) are color-coded based on LC integrity (median-split) and mean trajectories for subgroups are displayed (n = 33 younger adults are in the low and high LC group; n = 114 older adults are in the low and high LC group, respectively).

There were no statistically reliable age group differences in the association between memory performance (intercept, slope) and LC scores (likelihood ratio tests: all Δ*χ*^2^(Δ*df* = 1) **≤** 1.927, all *p* **≥** 0.165; see Supplementary Table 8). Thus, while we discovered a statistically reliable association between interindividual differences in memory performance and LC integrity only in older adults, we observed no reliable age group differences in this association (see Supplementary Table 9 for analyses across age groups).

Making use of the longitudinal nature of this data set (see Methods, section on study design and participants), we also examined associations between LC and memory performance using behavioral data that were obtained about 2.2 years before LC measurements were taken. We again observed that higher LC integrity was related to better memory performance among older adults, suggesting stable and lasting rather than short-lived LC–memory dependencies (see Supplementary Results 2.3.1.2).

### Lower age differences in rostral locus coeruleus intensity ratios relate to memory performance in older adults

Building upon reports of a spatially differentiated LC organization^21–23^, we investigated age differences in LC topography and their association with memory performance by analyzing LC data slice by slice. A non-parametric cluster-based permutation test^62^ revealed spatially heterogeneous age differences in LC ratios along the rostrocaudal extent of the nucleus (see Methods, section on analysis of age differences in the spatial distribution of locus coeruleus ratios). In comparison with younger adults, older adults showed a cluster of elevated intensity spanning caudal slices (65th–100th LC percentile; cluster permutation test: *p_corr_* < 0.001, [95% CI: < 0.001, < 0.001]) in line with neuromelanin accumulation across the life span^22, 42, 43^. In contrast, in rostral segments there was a trend towards decreased LC intensity in older adults (29th–41st LC percentile; cluster permutation test: *p_corr_* = 0.079, [95 % CI: 0.078, 0.080]; see Figure 5), in accordance with age-related decline in hippocampus-projecting LC segments^22, 23^. Age differences in LC ratios differ reliably across the rostrocaudal axis of the nucleus, as indicated by a significant Age group (Young/Old) × Topography (Rostral/Caudal) interaction (analysis of variance: *F*(1, 292) = 26.650, *p* < 0.001, *ηp*^2^ = 0.084, [95 % CI: 0.033, 0.149]^63^; see Figure 5).

**Figure 5.**
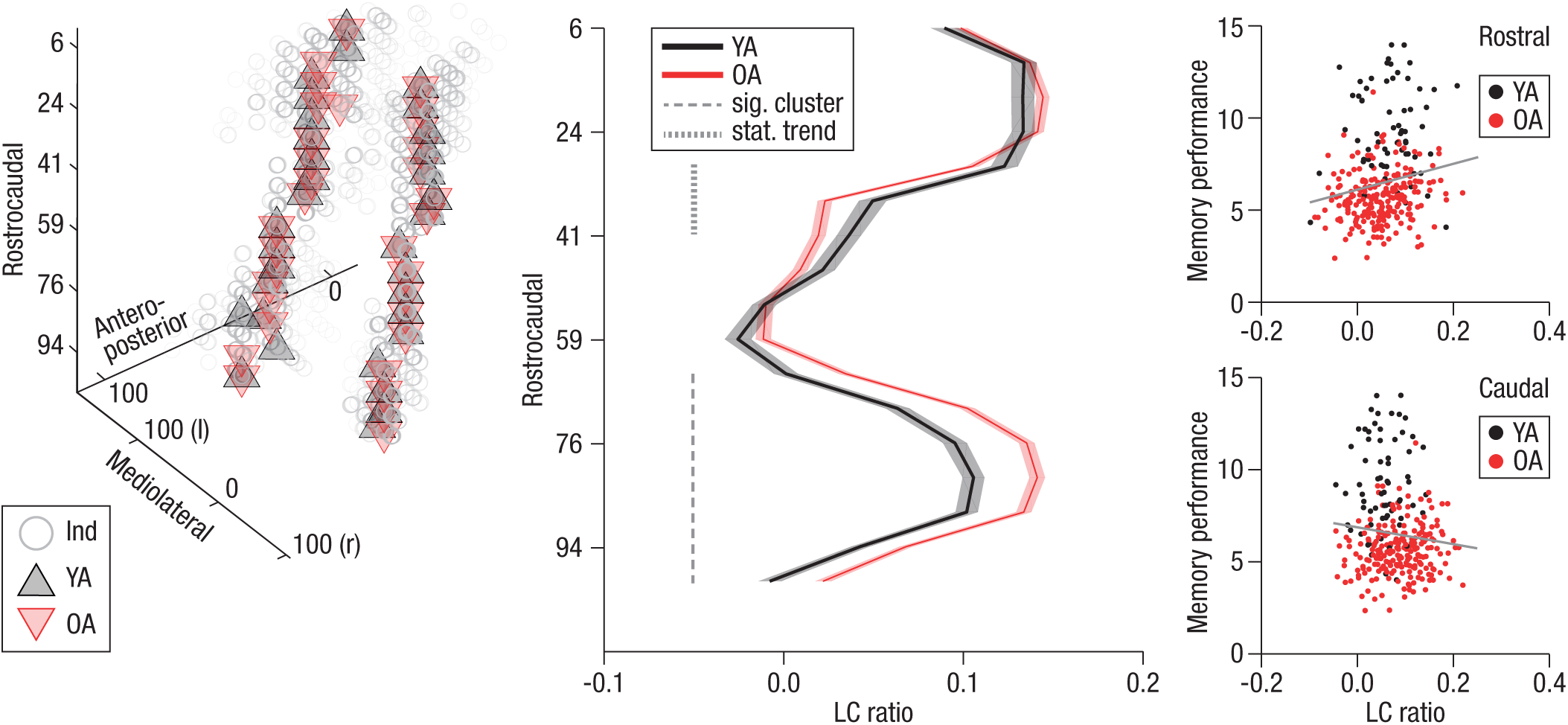
Topographical age differences in locus coeruleus intensity ratios and their functional significance. Left plot: On a group level, younger (YA; n = 66; black) and older adults (OA; n = 228; red) share highly congruent peak locus coeruleus (LC) locations. Grey circles indicate single subjects (Ind). Middle plot: A cluster-based permutation test revealed spatially confined age differences in LC intensity across the rostrocaudal axis. While older adults show significantly higher ratios in caudal segments (65th– 100th LC percentile; cluster permutation test: *p*corr < 0.001, [95% CI: < 0.001, < 0.001]), there is a trend for a reversed effect in rostral segments (29th–41st LC percentile; cluster permutation test: *p*corr = 0.079, [95 % CI: 0.078, 0.080]). Age differences in LC ratios differ reliably across the rostrocaudal axis of the nucleus, as indicated by a significant Age group (Young/Old) × Topography (Rostral/Caudal) interaction (ANOVA: *F*(1, 292) = 26.650, *p* < 0.001, *ηp*^2^ = 0.084, [90 % CI^63^: 0.040, 0.137]) Shaded lines represent ± 1 standard error of the mean (SEM; see Supplementary Figure 12 for the distribution of younger and older adults’ LC ratios along the rostrocaudal axis). Rostrocaudal, anteroposterior and mediolateral positions are expressed in percentiles relative to the total size of the observed LC. Right plots: The correlation between memory performance and LC ratios, calculated across YA and OA (n = 294), reaches statistical significance in the rostral cluster (*rs*(292) = 0.207, [95 % CI: 0.095, 0.314]*, p* < 0.001); it fails to reach statistical significance in the caudal cluster (*r_s_*(292) = –0.08, [95 % CI: –0.193, 0.035]*, p* = 0.172; difference between rostral and caudal correlations: *Z* = 3.385, *p*< 0.001).

To evaluate the functional significance of the observed topographical age differences, we related memory performance to LC ratios across all participants for each identified cluster. In caudal segments (65th–100th LC percentile), we observed no reliable association between LC ratios and initial recall performance (intercept; Spearman’s correlation: *rs*(292) = –0.08, [95 % CI: –0.193, 0.035], *p* = 0.172; see Figure 5). However, in rostral segments (29th–41st LC percentile), higher LC ratios were significantly associated with memory performance (Spearman’s correlation: *r_s_*(292) = 0.207, [95 % CI: 0.095, 0.314], *p* < 0.001; see Figure 5; difference between caudal and rostral correlations^64^: *Z* = 3.385, *p* < 0.001; for analyses within age groups, see Supplementary Table 11). Older adults with more youth-like intensity ratios in rostral LC segments also showed better memory performance. In sum, we observed a trend towards spatially confined age differences in LC ratios that were associated with memory performance and are in line with neurodegeneration of hippocampus- and forebrain-projecting LC segments^22^ in older adults.

## Discussion

Animal and post-mortem human studies suggest a link between memory performance in aging and the integrity of the central noradrenergic system^7, 10^. So far, in vivo research on humans, however, has been stymied by methodological difficulties in the reliable assessment of LC integrity^30, 40, 41^ (but see Hämmerer and colleagues^15^). Here, we took advantage of the paramagnetic properties of neuromelanin, a byproduct of NE synthesis, in T_1_-weighted MRI to image the LC in vivo^46, 48^. We assessed learning and memory performance in a large, healthy sample of younger and older adults along with structural MRI markers of LC integrity.

Our findings demonstrate reduced learning and memory performance in older relative to younger adults. Crucially, individual differences in learning and memory in a widely used neuropsychological test of memory functioning and, beyond that, across a variety of alternative memory tasks (see Supplementary Results 2.3.1.2), were positively related to LC integrity in older adults. Analyses making use of the longitudinal nature of this data set further indicate stable and lasting rather than short-lived LC–memory dependencies (see Supplementary Results 2.3.1.2). Moreover, we observed spatially confined age differences in LC intensity ratios. Older adults with more youth-like intensity ratios in rostral, hippocampus-projecting LC segments were better able to preserve memory performance. These results bridge a gap between animal and in vivo human research and provide important novel insights into the neural underpinnings of senescent cognitive decline in healthy aging.

We applied an iterative learning and memory task (RAVLT)^65^ that required subjects to encode, consolidate, and retrieve verbal information and thus captures the dynamic nature of memory^53^. Our analyses allow the study of two psychologically distinct factors, namely, initial recall (i.e., performance after the first learning trial, corresponding to a standard one-trial memory assessment) and learning (i.e., changes in performance with practice). Age differences were mainly observed in initial recall performance (see Supplementary Results 2.1.3 and Supplementary Discussion).

Neurodegenerative changes have been suggested to be determinants of interindividual differences in learning and memory performance in late life^51, 66^. Here, we investigated the role of the LC, one of the first targets of neurodegenerative diseases such as Alzheimer’s and Parkinson’s, in senescent memory decline^8, 9, 11^. In particular, we exploited the paramagnetic properties of neuromelanin-metal compounds in T_1_-weighted MRI to index the integrity of the LC in vivo^46^ (see ^67, 68^ for recent discussions of LC contrast mechanisms). Neuromelanin accumulates non-linearly in LC cells across life with a peak concentration at around 50–60 years of age and subsequently stablizes^38^ or declines, probably due to preferential loss of pigmented cells^42, 43^. Thus, neuromelanin forms a reliable natural contrast agent, particularly in older adults, that can be harnessed in cell count^18^ and MRI studies^48^ to index LC integrity. However, neuromelanin may be a less effective indicator of LC structure in younger adults due to the many LC neurons that do not yet contain neuromelanin, based on post-mortem analyses using tyrosine hydroxylase (i.e., the rate-limiting enzyme in NE synthesis) as a marker of LC neurons^22^. From the fifth decade onwards the numbers of LC neurons identified by tyrosine hydroxylase and neuromelanin markers were similar^22^. We successfully established a semi-automatic procedure that extracted individual LC intensity ratios across the rostrocaudal extent from high-resolution neuromelanin-sensitive brainstem MRI in younger and older adults. In particular, we pooled over aligned scans to facilitate LC delineation at a group level^61^, which in turn was used to restrict automatized extraction of individual peak intensities^68, 69^. The LC volume of interest (search space) generated at the group level matches recent histological analyses that reported a dense packing of noradrenergic cells in a thin central LC compartment and dispersion towards rostral and especially caudal extremities^18^. We demonstrated high validity of the proposed semi-automatic procedure by comparisons to both previously published LC maps^41, 61^ and manual LC intensity ratings (see Supplementary Results 2.2.1.1–2). Repeated measurements of an independent younger adult sample further confirmed high reproducibility of the intensity assessment (see Supplementary Results 2.2.1.3). However, as reported earlier for neuromelanin-sensitive sequences, our analyses do not capture the complete LC extent^47, 70^ leading to lower volume estimates than those obtained post-mortem studies^19^. In their histological analyses, Fernandes and colleagues describe a common, central LC zone which is shared across all subjects^18^. Accordingly, the segments with the most dispersed noradrenergic cells in the caudal LC show less overlap on a group level and are thus arguably more difficult to detect using a semi-automatic approach with a conservative intensity threshold (i.e., 4*SD* above mean intensity of a reference region^68, 69^). This circumstance is likely exacerbated by partial volume effects due to the relatively thick slices in most neuromelanin-sensitive sequences^70^. However, manual^61^ and threshold-free automatic approaches^41^ also appear to be affected by this challenge as evident from their low correspondence in caudal slices (MNI Z ≤ –30: 6.91 % of Keren et al. in Betts et al., 4.63 % vice versa^41, 61^; see Figure 2d). We thus refrained from attempts to capture the most caudal LC segments via manual segmentation, a lowering of intensity thresholds, or both, in order to effectively separate LC from the nuclei subcoerulei^17^.

To investigate the link between LC integrity and memory in aging humans, we estimated latent LC integrity scores using structural equation modeling. Here, we integrated over slices and hemispheres to derive a single measure reflecting LC integrity. Compatible with spatially confined age differences^22, 41, 61^, differences between younger and older adults in average LC scores failed to reach statistical significance. In older adults, higher initial recall performance and steeper learning curves were found in subjects with high LC integrity. Even when integrating over a variety of tasks, our findings corroborate a positive dependency of memory performance on LC integrity (see Supplementary Results 2.3.1.2), in line with a general mechanism beyond specific mnemonic domains (e.g., emotional memory^15^).

Besides regulating healthy cognition^21, 26, 29, 31, 32^ the LC subserves critical neuroprotective functions^11, 71, 72^. For instance, NE protects against neuroinflammation via the regulation of inflammatory gene expression and directly enhances clearance of aggregated ß-amyloid via activation of microglia^11, 72^, two major threats to the aging brain^73^. This led Robertson and colleagues^74^ to suggest that repeated activation of the LC-NE system (e.g., by exposure to novelty^37^) helps maintain cognitive functionality in aging despite underlying neuropathological changes. At the same time, the LC is susceptible to neurodegeneration in aging. Its vulnerable anatomical location next to the fourth ventricle exposes it to toxins and its high energy demand increases the risk of oxidative stress over time^71^. Thus, while in aging LC-NE’s neuroprotective function is especially important, the loss of noradrenergic cells may have wide-ranging consequences for both cognition and brain health. Whereas we observed a positive association between LC integrity and memory performance in older adults, we did not find a robust relation in younger adults. However, the association between LC integrity and memory did not differ reliably between groups. Thus, while we discovered an association between interindividual differences in memory performance and LC integrity in older but not younger adults, we did not observe reliable group differences which may hint to comparable associations independent of age. This appears to contrast with the notion of an increasingly important role of the LC-NE system in aging^11^. However, to constitute a reliable proxy for neuronal density within the LC^48^ neuromelanin-sensitive MRI requires a sufficient neuromelanin-saturation of the nucleus. Post-mortem studies indicate that neuromelanin concentration within the LC peaks around late middle adulthood (∼ 50 years of age^43, 75^). Thus, the lower sensitivity of neuromelanin as a LC integrity proxy in younger adults may have impeded finding reliable associations within this subgroup^15, 76^ as well as differences between age groups. We observed spatially confined age differences in LC intensity ratios^41, 61^. In older relative to younger adults, caudal LC sections showed significantly elevated intensity ratios in line with neuromelanin accumulation across the life span^42, 43^. In contrast, there was an absence of increased signal in rostral LC and, descriptively, even a tendency towards reduced intensity ratios in older adults. This may suggest neurodegeneration of rostral LC segments that are densely connected to key memory structures like the forebrain and hippocampus^21–23^. Consistent with this possibility, LC intensity ratios in rostral but not caudal segments were positively associated with memory performance. This finding is in line with a series of cell counting studies that point towards specific loss of rostral LC compartments even in healthy aging (as covered by a review of early studies^22^, but as recently discussed^11^ there are also recent observations^16, 18, 38, 39^ of stable LC cell counts in healthy aging).

At least three well-documented mechanisms of noradrenergic action may explain the observed general memory-promoting effect of high LC integrity. First, NE release in the sensory cortices increases signal-to-noise ratios by silencing spontaneous activity while sparing or even facilitating stimulus-evoked responses^21, 32^, presumably in self-enhancing feedback loops with glutamate^29^. In addition, NE improves the temporal organization of neuronal responses to sensory stimulation (i.e., spike rhythmicity) and thus increases perceptual acuity^32, 77, 78^. Note that noisier information processing has been linked to deficient neuromodulation with effects on higher-order cognitive functions like attention and working memory^79^. Second, via action on post-synaptic α2A-adrenoceptors in the prefrontal cortex, moderate levels of NE facilitate delay-related activity, which is considered a cellular analogue of working memory^26^. Accordingly, pharmacological manipulation of NE levels in aged monkeys ameliorated attention and working memory deficits^14^. Hence, LC integrity may promote attentional and control mechanisms implicated in successful episodic memory performance^80^. Third, in the amygdala and hippocampus, NE modulates synaptic strength and facilitates synaptic plasticity^31, 32^. More specifically, via its action on ß-adrenoceptors, the LC modulates LTP and LTD, major determinants of long-term memory^33, 34^. In sum, NE released from the LC alters perception, attention and memory at multiple cortical and subcortical sides, crucially implicating LC integrity in learning and memory in aging.

Recent analyses of large postmortem samples have indicated that the LC is among the earliest brain regions to show hyperphosphorylated tau, a hallmark of Alzheimer’s disease, which accumulates in the nucleus linearly with the progression of the disease^8, 16, 81^ (but see^82^ for a different observation and^83^ for a commentary). In tauopathies like Alzheimer’s, misfolded tau proteins are thought to spread in a stereotypical, transcellular propagation pattern through neural networks^8, 82^ which may lead to tau pathology in prominent noradrenergic projection targets such as the medial temporal lobe^72^. Elevated tau levels are associated with more aggregated amyloid-ß, the second hallmark of Alzheimer’s disease, and worse cognition^73^. A mere accumulation of subcortical tau could be harmless without evidence for neurodegenerative consequences. However, LC volume decreases by about 8% per Braak stage increment (i.e., a classification system that allows staging the progression of tau spread) followed by death of LC cells (from middle Braak stages on^16, 84^) that correlates with cognitive decline (for a review on the impact of hyperphosphorylated tau on LC, see^72^). Of note, LC cell loss in Alzheimer’s disease reaches profound levels (∼50–80 %^71^) and even surpasses neurodegeneration of the cholinergic nucleus basalis^85^. In animal models of Alzheimer’s disease, LC lesions exacerbate both neural and behavioral decline^71^. In particular, in a mouse model of tauopathology, LC-ablated mice showed impaired memory and increased neuronal loss in the hippocampal formation compared with non-ablated animals^72, 81, 82^. Further, in a rat model with both ß-amyloid and tau pathology, chemogenic activation of the LC rescued hippocampus-dependent behavior^84, 88^. Together, this indicates that in Alzheimer’s disease LC neurodegeneration acts synergistically with tau pathology to drive pathological changes and does not merely constitute an epiphenomenon^87^. The threshold between healthy aging and early manifestations of neurodegenerative diseases like Alzheimer’s is yet to be defined^84^. Most likely, both LC neurodegeneration and the transition from healthy cognition in aging to pathologic conditions (i.e., mild cognitive impairment, dementia) occur along a continuum^76^. While all but two of the older adults tested in this study scored above-threshold on a dementia screening (i.e., Mini Mental State Examination^89^) and demonstrated a high level of functionality, neither our cognitive nor neural data can rule out the existence of early levels of neuropathology (e.g., subcortical accumulation of pretangle tau^8^). We thus cannot rule out that both normal aging and neuropathologic changes affected locus coeruleus integrity in our older adult sample.

The present results underscore the utility of non-invasive, in vivo markers of LC integrity as indicators for preserved memory performance in human aging and extend our knowledge about the role of the LC-NE system in senescent decline. Using non-invasive in vivo MRI, we discovered spatially confined and functionally relevant differences between younger and older adults in those segments of LC that are connected to key memory structures like the hippocampus. Importantly, structural equation modeling revealed reliable and stable positive associations between LC integrity and general episodic memory among older adults. We conclude that older adults with more preserved LC integrity are equipped with more proficient episodic memory. This finding, which needs to be corroborated by long-term longitudinal evidence, adds specificity to the general proposition that brain maintenance is a key feature of successful cognitive aging^1^.

## Methods

### Study design and participants

Data was collected within the framework of the Berlin Aging Study-II (BASE-II^90–92)^. BASE-II is a multi-disciplinary and multi-institutional longitudinal study that investigates the cognitive^93^, neural, physical, and social conditions that are associated with successful aging. For an extensive study description, please refer to the study’s website (https://www.base2.mpg.de/en) and papers^90–92^.

Cognitive and neuro-imaging data of younger and older participants were collected at the Max Planck Institute for Human Development (Berlin, Germany) at two time points (T1 and T2) between 2013 and 2016. On average, data acquisitions were 2.21 years apart (standard deviation (*SD*) = 0.52, range = 0.9–3.2). Cognitive and neuro-imaging data were collected on separate occasions at each time point (mean interval = 9.16 days, *SD* = 6.32, range = –2–44; for T2). Locus coeruleus (LC) data were acquired only at the T2

A subset of 323 BASE-II participants underwent magnetic resonance imaging (MRI), with 24 of these excluded before analysis due to missing or incomplete neural (n = 19) or cognitive (n = 5) data. After visual inspection of brainstem MRI, five additional participants (0 female; mean age: 76.66 years; *SD* = 1.64; range = 74.93–78.74; at T2) were excluded from further analyses due to excessive movement artifacts (n = 2) or incorrect scan positioning (n = 3). The final MRI subsample (see Supplementary Table 1) included 66 younger adults (22 female) with a mean age of 32.5 years (SD = 3.53, range = 25.41–39.84; at T2) and 228 older adults (82 female) with a mean age of 72.29 years (*SD* = 4.11; range = 62.53–83.16; at T2). All participants with full cognitive and MRI data were included in the study, i.e., we did not conduct a formal power calculation, given that there was no available prior evidence on the studied phenomena. Neurological and psychiatric disorders, a history of head injuries, or intake of memory-affecting medication precluded inclusion in the study. All eligible participants were MRI-compatible, right-handed, and had normal or corrected-to-normal vision.

The cognitive and MRI assessment were approved by the Ethics Committees of the Max Planck Institute for Human Development and the German Psychological Society (DGPs), respectively. Participants signed written informed consent and received monetary compensation for their participation. All experiments were performed in accordance with relevant guidelines and regulations.

### Cognitive data assessment

The baseline cognitive assessment of BASE-II (T1) and its follow-up (T2) included an assessment of working memory, episodic memory and fluid intelligence in small group sessions of about 6 individuals using a computerized battery of 21 tasks (for a description of the complete battery, please refer to^93^). Each test session lasted approximately 3.5 hours. At T1, a second cognitive assessment was scheduled one week after the first session to test for long delayed recall performance.

Our hypotheses focused on the Rey Auditory Verbal Learning Test (RAVLT), a standardized and validated neuropsychological tool that provides information about participants’ learning and delayed recall performance^65^ (see Figure 1): Subjects first learned a 15-word list that was auditorily presented via headphones. The task was composed of five learning trials each followed by a free recall period in which participants entered the words they remembered via keyboard (trial 1–5; recall of learning list). After initial learning, a 15-word distractor list, containing semantically unrelated words was presented, followed by a free recall phase (trial 6; recall of the distractor items). Next, participants were again asked to freely recall only items presented in the initial list (trial 7; recall of learning list after distraction). Another free recall test was administered after a delay of 30 minutes (trial 8; delayed recall of learning list). The verbal learning memory task ended with a recognition memory test that included 50 items (learning list: 15, distractor list: 15, similar lures: 20). At T1, participants were re-invited 7 days later for a final long delayed free recall test (trial 9; long delayed recall of learning list). After the initial learning cycles, the correct word list was never re-presented to subjects. The same word list was used at T1 and T2.

Making use of the comprehensive cognitive battery available for this data set, we additionally integrated performance over a variety of episodic memory tasks to retrieve a general measure of episodic memory as previously described^93^ while explicitly excluding RAVLT data (see Supplementary Results 2.1.4). In particular, we incorporated information of a visual face– profession, object–location, and scene encoding task. For detailed task descriptions, please refer to^93^. In short, the visual face–profession task (FPT) involved subjects studying the 45 pairs of face images and profession words. The tasks consisted of an incidental encoding phase, a 2-minute distraction phase, and finally a recognition memory task including old, new as well as rearranged face–profession pairs. Recognition memory (hits – false alarms) for rearranged pairs was used as the performance index. In the visual object–location memory task (OLT), participants encoded the location of 12 digital photographs of real life objects on a 6 x 6 grid. After encoding, objects reappeared next to the grid and subjects were asked to reproduce the correct location by placing the items in the correct grid position. The sum of correct placements was used as the performance index. Finally, in a visual scene encoding task (IOST), participants incidentally encoded 88 scene images by performing indoor/outdoor judgments on each image. The encoding phase was followed by an old/new recognition memory test which included confidence judgments. Recognition memory (hits – false alarms) was tested after a delay of approximately 2.5 hours and served as the performance index.

### Magnetic resonance imaging data assessment

Structural and functional MRI data were collected on both time points (T1, T2) employing a 3-Tesla Siemens Magnetom Tim Trio Scanner with a standard 12-channel head coil. Only those sequences used in the current analyses are described below.

A three-dimensional T_1_-weighted magnetization prepared gradient-echo (MPRAGE) sequence with a duration of 9.2 min and the following parameters was applied: TR = 2500 ms, TE = 4.770 ms, TI = 1100 ms, flip angle = 7 °, bandwidth = 140 Hz/pixel, acquisition matrix = 256 × 256 × 192, isometric voxel size = 1 mm^3^. Pre-scan normalize and 3D distortion correction options were enabled.

Based on this whole-brain MPRAGE sequence, a neuromelanin-sensitive high-resolution, two-dimensional T_1_-weighted turbo-spin echo (TSE) sequence was aligned perpendicularly to the plane of the respective participant’s brainstem. Acquisition of the TSE sequence took 2 × 5.9 min, and the following parameters were used: TR = 600 ms, TE = 11ms, flip angle = 120 °, bandwidth = 287 Hz/pixel, acquisition matrix = 350 × 263 × 350, voxel size = 0.5 × 0.5 × 2.5 mm^3^. Each TSE scan consisted of 10 axial slices with a gap of 20 % between slices, which covered the whole extent of the pons. Pre-scan normalize and elliptical filter options were enabled. The TSE sequence yielded two brainstem MRIs per participant, each resulting from 4 (online) averages.

For some participants, specific absorption rate (SAR) limits were exceeded, which precluded further scanning with the current set of parameters (cf.^47, 68^). To avoid data loss while maintaining contrast settings, for those participants the maximal number of TSE slices was reduced. Thus, our sample consists of the following slice distribution: 243 × 10 slices, 38 × 9 slices, 27 × 8 slices, 6 × 7 slices, 1 × 6 slices, 3 × 11 slices (acquired before adjusting the number of slices in the sequence protocol). TSE scans are available only for T2.

### Cognitive data analysis

We applied structural equation modeling (SEM) to analyze inter- and intra-individual differences in verbal learning and memory performance using the Ωnyx 1.0-991 software package^94^ with full information maximum likelihood estimation (FIML). SEM offers a multivariate approach in which repeatedly observed (manifest) variables can be used to examine hypotheses about unobserved (latent) variables. Latent variables have the benefit of accounting for measurement error in observed scores and thereby increasing statistical power^57, 58^.

We assessed the adequacy of the proposed growth models (i.e., a specific variant of SEM) by testing for differences between the model-implied and empirically observed covariance matrices^60^. The *χ*^2^-test formally tests for equity of the covariance matrices. However, it is particularly sensitive to sample size and should be interpreted with caution in large samples^59, 60^. Thus, we examined two additional frequently reported fit indices (RMSEA, CFI). In contrast to the *χ*^2^-test, the CFI is not influenced by sample size. Finally, the adequacy of the quadratic growth model was tested against nested competing alternative models (e.g., intercept only and linear growth models) using likelihood ratio difference tests^57, 60^. In the case of non-nested competing alternative models, we compared the Akaike Information Criterion (AIC), favoring the model providing the smallest AIC^57^.

After establishing model fit, age differences in parameters of interest (e.g., means and (co)variances of intercept and slope factors) were tested by fixing parameters to equity between groups and comparing model fit to a model in which parameters were freely estimated using a likelihood ratio difference test ^57, 60^ (see Results, section on initial recall performance is lower in older adults). In addition, for all models, parameters of interest are listed with their point estimate and the confidence interval which is based on the standard error (SE) of the estimate (i.e., point estimate ± SE × 1.96), computed by the inverse of the Hessian^94^. The confidence interval allows a quick indication of whether parameters differ reliably from zero. Note, however, that the estimation of confidence intervals focuses on a single parameter and does not take the correlation with the remaining parameters into account, thus leading to lower statistical power^94^. Statistical inferences are therefore based on likelihood ratio tests as previously suggested^57, 58, 94^. When applicable, we also added standardized parameter estimates (i.e., correlation coefficients for instance for associations between latent LC and memory estimates).

In a similar vein, we integrated performance over a variety of memory tasks to estimate a general episodic memory score using a multiple-group single factor structural equation model as previously described^93^(see Supplementary Figure 4). In particular, in each age group performance in the visual face – profession, object – location, and scene encoding tasks served as manifest variables and loaded on a single latent episodic memory factor. Factor loadings (other than the first, which was fixed to one) were estimated freely but were constrained to be equal across groups. Adequacy of the proposed model was assessed using a *χ*^2^-test as well as two additional fit indices (RMSEA, CFI).

### Magnetic resonance imaging data analysis

We established a multi-step analysis procedure to extract individual LC intensity values that addressed a series of methodological concerns raised in previous research: First, TSE scans of the LC typically demonstrate a low signal-to-noise ratio due to the high spatial resolution needed to image the small nucleus^68^. In addition, the LC shows amorphous borders, complicating the manual segmentation of the structure^41, 70^. Thus, the use of semi-automatic threshold-based approaches was recently suggested to circumvent manual tracing of the nucleus and associated errors as well as potential biases (e.g., non-blinded raters^47, 68, 69^). Second, a reliable coregistration of brainstem MRI to standard space poses a major methodological challenge with standard software packages that can be addressed by the use of study-specific templates and coregistration masks^41^. Third, age- and disease-related alterations of LC integrity are not homogeneous across the rostrocaudal extent of the structure^9, 41, 61^. Increased specificity and sensitivity to detect such conditions may be reached by assessing intensity across the whole rostrocaudal axis^47^. Fourth, due to the use of multiple measurements and online averaging, the acquisition time for most neuromelanin-sensitive sequences exceeds 10 minutes, which increases the risk of head motion artifacts and thus reduced data quality^68^. To explicitly correct for this, images from each measurement can be saved separately and averaged after application of motion-correction methods^68^.

In the implemented analysis procedure (see Supplementary Figure 1), we first pooled across subjects over aligned TSE scans to increase signal-to-noise ratio and facilitate LC delineation using a template-based approach (cf.^41, 61^; Steps 1–3). On a group level, LC location was determined semi-automatically (cf. ^68, 69^) to restrict fully-automatized extraction of individual peak intensity values across the rostrocaudal extent (Steps 4–5). For each participant, data from two scans (both from T2) were assessed, and extracted values were averaged to obtain a more reliable estimate of LC intensity.

### Coregistration and standardization of magnetic resonance imaging data

Prior to analysis, all whole-brain and brainstem MRI scans (MPRAGE, TSE) were up-sampled to twice the initial matrix size by sinc-interpolation to improve visualization of the LC as previously described (cf.^61^).

Initially (Step 1), using Advanced Normalization Tools v. 2.1 (ANTS^95, 96^), a group whole-brain template (Template_whole_) was generated from all available MPRAGE scans from T2. In this iterative process, (a) individual native space scans (MPRAGE_native_) were coregistered to a common group or template space and (b) coregistered scans (MPRAGE_template_) were then averaged to form the group whole-brain template. An average of all input files served as an initial fixed image. One linear (rigid, then affine) registration of input files was followed by six non-linear registration iterations. Each non-linear registration was performed over three increasingly fine-grained resolutions (30 × 90 × 20 iterations). We applied a N4 bias field correction on moving images before each registration. Cross-correlation (CC) was used as the similarity metric and greedy Symmetric Normalization (SyN) as transformation model for non-linear registrations^97^. Template update steps were set to 0.25 mm.

Next (Step 2), within each subject, we coregistered the native space brainstem scan (TSE_native_) to the whole-brain scan that had been moved to template space (i.e., MPRAGE_template_; see Step 1). For this, we first performed linear (rigid, then affine) followed by nonlinear (SyN) registration steps^97^. Histogram matching was enabled since we registered scans from the same modality (T_1_-weighted). This intermediate within-subject coregistration was implemented to bring individual brainstem scans to a common (whole-brain) space and thereby correct for differences in acquisition angle that would reduce precision of the template building (see Step 3).

Finally (Step 3), the aligned brainstem scans were fed into a second template building run, resulting in (a) individual brainstem scans coregistered to template space (TSE_template_) and (b) a group brainstem template image (Template_slab_). The same parameters were used were used for both template building runs (Step 1, Step 3). The linear and non-linear transformation matrices of the intermediate within-subject coregistration (Step 2) and template building (Step 3) were concatenated and applied to the native space brainstem images (TSE_native_) to achieve spatial standardization (TSEtemplate) within a single transformation step (cf.^61^). To correct for minor deviations between whole-brain and brainstem template space, we coregistered the two group templates using a linear (rigid, then affine) followed by nonlinear (SyN) registration. Since we registered a whole-brain image and a slab here, a coregistration (or “lesion”) mask was generated that restricted registration to the brainstem, i.e. voxels falling inside the Template_slab_ (cf.^41^).

### Semi-automatic locus coeruleus segmentation and intensity assessment

Compared to individual scans, the group brainstem template demonstrated a high signal-to-noise ratio and markedly increased intensities in left and right brainstem sections next to the fourth ventricle corresponding to hypothesized LC locations (see Figure 3b). In order to circumvent manual segmentation on the brainstem template, we thresholded the image (Step 4) allowing only the brightest voxels to remain visible using a custom Matlab function (cf.^68, 69^; Mathworks Inc., Sherborn, MA, USA, including the SPM software toolkit^98^). In particular, we applied a data driven approach using a threshold based on the intensity in a dorsal pontine reference region (see below) that effectively isolated patches of high image intensity next to the fourth ventricle. The threshold was calculated for each template slice that covered the full brainstem (n = 31), separately for each hemisphere, as follows:

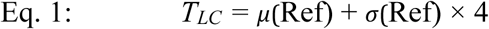

Whereby *μ* and *σ* specify the mean and standard deviation of the intensity in the pontine reference (Ref) region, respectively. *T_LC_* denotes the resulting intensity threshold. The cut-off of 4 standard deviations above the mean reference intensity was selected on the basis of previous research^68, 69^ and was confirmed visually as generating a volume that matched prior mapping studies of the LC^41, 61^. A lateralized threshold was computed for each slice to counteract previously reported unilateral biases in brainstem signal intensity^61, 70^ and intensity fluctuations along the rostrocaudal axis. Next, we generated a volume of interest (VOI), subsequently referred to as search space, that included only those areas that survived thresholding. The resulting bilateral LC search space covered 17 slices and included 409 voxels in total (see Figure 3c). In addition to the LC search space, we also defined a larger rectangular dorsal pontine reference search space, spanning 35 × 10 voxels on all 17 slices on which the LC remained visible (5950 voxels). The bilateral LC and reference search spaces were then split at midline to allow for bilateral and lateralized analyses.

To obtain LC intensity ratios for all subjects (Step 5), we masked individual brainstem scans in template space (i.e., TSE_template_) with the binarized LC search space. This was first done with a bilateral, followed by unilateral (left, right) search spaces. We thereby restricted the area from which intensity values could be extracted to potential LC coordinates. We then automatically selected the peak (maximal) intensity voxel within each masked scan for each slice and stored the corresponding intensity value and x/y/z coordinates^41^. In the same vein, we masked individual scans with a binarized reference search space and extracted intensity values and coordinates. LC intensity ratios were then computed for each slice using the following formula^47^:

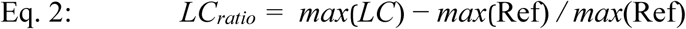

Whereby LC denotes the intensity of the locus coeruleus and Ref indicates the intensity of the pontine reference region. Finally, for each hemisphere we computed a LC ratio across all slices, which was used to probe LC–cognition associations in a SEM framework (i.e., we calculated a ratio score of the peak LC intensity relative to the peak reference intensity).

### Comparison to previously published locus coeruleus masks

In order to judge the validity of the generated search space, we plotted the retrieved peak intensity coordinates for each subject’s left and right LC^41^. The aggregated maximum intensity plot was then converted to express the likelihood to find the LC for any given voxel (i.e., a LC location probability map; cf.^41, 70^). We thresholded the image to show only locations that were shared across participants (i.e., the upper 90 percentiles) and spatially standardized the probability map in a two-step procedure. First, we coregistered the whole-brain template (see Step 1) to 0.5 mm iso-voxel MNI space using linear (rigid, then affine), followed by nonlinear (SyN) registration. Since the brainstem template was registered with the whole-brain template (see Step 3), in a second step we were able to apply the transformation matrices (Template_slab_ → Template_whole_ and Template_whole_ → MNI_0.5_) to the LC probability map. In standard space, we calculated the overlap between the probability map and previously published LC masks^41, 61^; see Figure 3d and Supplementary Results 2.2.1.1). To facilitate comparability of study results, we freely share the resulting LC probability map with the neuroscientific community (https://www.mpib-berlin.mpg.de/LC-Map).

### Comparison to manually assessed locus coeruleus intensity

Almost all published MRI studies that indexed LC integrity used a manual procedure to assess intensity values (for a recent review, see^47^). To demonstrate the validity of our approach, two independent, blinded raters (research assistants) manually assessed LC and reference intensity values for all subjects using a standard procedure^75^. On each axial slice demonstrating increased intensity in anatomically probable LC locations, raters placed three non-overlapping quadratic 1 mm² regions of interest (ROIs) in areas in which signal was most evident (cf.^75^). Raters were instructed not to place the ROIs directly adjacent to the fourth ventricle to avoid partial volume effects contaminating the signal (cf.^76^). To measure reference intensity, three rectangular non-overlapping 10 mm² ROIs were placed on the same slices in the dorsal pons, centered medially between the left and right LC ROIs (cf.^75^; see Supplementary Figure 2). Raters additionally indicated the slice with peak signal intensity. Image processing was conducted using the image j software (https://imagej.nih.gov/) in a room with constant illumination. Ratings were performed on offline-averaged, non-interpolated brainstem scans in native space. For four subjects manual assessments were not available from both raters and those subjects were thus dropped from the evaluation. For analyses, the peak intensity for each slice was first determined for LC (left, right) and pons across ROIs, and then the peak intensities were selected across slices. We calculated the consistency of raters using intraclass correlation coefficients (ICC; two-way mixed model with absolute agreement, using IBM SPSS Statistics v. 24) and then, after demonstrating high accordance, averaged ratings across hemispheres and raters to obtain a single stable intensity estimate per subject (see Supplementary Results 2.2.1.2). Comparisons between manually and automatically assessed LC intensity values (see Step 5) were eventually assessed by means of ICC. For further analyses, we also calculated LC intensity ratios for both hemispheres using Eq. 2 (cf. semi-automatic LC intensity ratio calculation; see Supplementary Results 2.2.1.2 and 2.5).

### Reproducibility of semi-automatically assessed locus coeruleus intensity ratios

In order to judge the temporal stability of the semi-automatic method, we repeatedly scanned a small number of younger adults who were not participating in the study over the course of 1–2 weeks (n = 3; 2 female; mean age: 23.667 years; *SD* = 1.578 years). Subjects were scanned four times resulting in eight brainstem images per participant (see MRI data assessment; 22 scans in total; missing data for n = 1 at the last time point). Data was sinc-interpolated and fed into another template-building procedure as described above (see Steps 1–3). The resulting brainstem template (template_slab_reproducibility_) was coregistered to the study’s brainstem template (template_slab_) using linear (rigid, then affine) followed by nonlinear (SyN) registration. This process effectively enabled alignment of each participant’s scans with the group brainstem template and thus allowed us to use the established search spaces to compute LC intensity ratios as described above. Finally, ICC were calculated for averaged left and right LC ratios to evaluate the stability of the proposed method. Note that we analyzed intensity ratios here since we compared scans acquired at different time points that are subject to overall scanner fluctuations.

### Comparison to semi-automatically assessed locus coeruleus intensity in native space

We additionally extracted individual LC and reference intensity from native space brainstem scans (TSE_native_) to confirm that intensity values retrieved from template space (TSE_template_) are not an artifact of coregistration and template building (Steps 1–3). For this, we used the concatenated inverse transformation matrices of Steps 2 and 3 to project the search spaces back to individual subjects’ coordinates. In order to compare spatial locations of native space values across subjects, however, a common frame of reference was necessary (cf.^61^). Therefore, before warping, we split the LC and reference search spaces into segments. Each but the most caudal segment contained three slices, reflecting the resolution difference between native and template space (i.e., 1 × TSE_native_ : 3 × TSE_template_; 1.5 mm : 0.5 mm). The six resulting segments were separately transformed to native space. We then masked native space scans (TSE_native_) with binarized LC and reference (segment) search spaces and assessed peak intensities as described above (cf. Step 5). The two native space brainstem images of each participant (see MRI data assessment) were linearly and non-linearly coregistered before masking. To judge the correspondence between automatically assessed LC intensity values from native and template space, we calculated ICC (see Supplementary Results 2.2.1.4). For further analyses of native space values, we computed LC intensity ratios for both hemispheres as described above using Eq. 2 (see Supplementary Results 2.4).

### Analysis of age differences in the spatial distribution of locus coeruleus ratios

To investigate age differences in LC intensity ratios along the rostrocaudal extent of the nucleus^9, 22, 41, 61^, we calculated non-parametric, cluster-based, random permutation tests as implemented in the Fieldtrip toolbox. These effectively control the false alarm rate in case of multiple testing^62^. In short, first a two-sided, independent samples *t*-test is calculated for each slice. Neighboring slices with a *p*-value below 0.05 were grouped with spatially adjacent slices to form a cluster. The sum of all *t*-values within a cluster formed the respective test statistic. A reference distribution for the summed cluster-level *t*-values was computed via the Monte Carlo method. Specifically, in each of 100,000 repetitions, group membership was randomly assigned, a *t*-test computed, and the *t*-value summed for each cluster. Observed clusters whose test statistic exceeded the 97.5th percentile for its respective reference probability distribution were considered significant^62^. Cluster permutation tests were calculated first across hemispheres and then separately for each hemisphere on LC ratios assessed from template (TSE_template_) and native space (TSE_native_; see Supplementary Results 2.2.2 and 2.4.1). Here, we used a two-sided test with a significance level (α) of 0.025. In the following, we thus report the doubled cluster *p*-values to facilitate readability (i.e., *p* ≤ 0.05 is considered significant for all statistics). Reliable clusters were followed up means of a mixed-effects analysis of variance (ANOVA) with the factors Age (YA; OA) and Topography (Rostral/Caudal) to evaluate spatially confined age differences in LC ratios. For this, LC intensity ratios were averaged within subjects within the ranges of the observed clusters.

To judge the functional significance of identified topographical age differences, we investigated the association between memory performance and LC ratios in the obtained clusters, first, irrespective of age group and then within each group. In particular, for each cluster, the relation between initial recall performance (intercept) and LC ratios (average over slices within the identified cluster) was assessed by means of Spearman’s rank correlations.

Finally, to examine inter-hemispheric differences in LC ratios^61, 70^, we additionally computed related-samples Wilcoxon signed rank tests. We first calculated a statistic across age groups, which we then followed up by analyses within each age group.

### Estimation of latent locus coeruleus integrity scores

As done for the cognitive data, we applied SEM to analyze interindividual differences in LC ratios. We generated a multiple-group model including average LC ratios of each hemisphere as observed variables (see Step 5; see Supplementary Figure 6). In each group, the two observed variables loaded on a single latent LC integrity factor. Factor loadings (other than the first, which was fixed to one) were estimated freely but were constrained to be equal across groups. Adequacy of the proposed model was assessed using a *χ*^2^-test as well as two additional fit indices (RMSEA, CFI). The analyses were first completed with automatically assessed LC ratios (TSE_template_). To demonstrate the stability of the findings, we repeated the same steps with values assessed in native space (TSE_native_) and manually assessed intensity ratios. Qualitatively similar results were obtained (see Supplementary Results 2.5.1.1).

### Analysis of associations between locus coeruleus integrity scores and memory performance

After generating structural equation models for our cognitive and neural measures, respectively, we set out to link these modalities. That is, we were interested in assessing the relation between interindividual as well as intra-individual differences in verbal learning and memory performance and interindividual differences in LC scores. For this, we first built a unified model merging the verbal learning and LC models described above for each time point (see Figure 2). We investigated associations between our performance and LC measures by allowing for freely estimated covariances on a latent level (shown in black, Figure 2). Models were estimated using LC ratios assessed in template space (TSE_template_). To assess the stability of the obtained findings, the same analyses were repeated with LC ratios assessed in native space (TSE_native_) and manually assessed LC ratios (see above). Qualitatively similar results were derived (see Supplementary Results 2.5.1.2). In order to evaluate the generalizability of the expected LC–memory association, we repeated the analyses, this time merging the neural SEM with the general episodic memory SEM (see Methods, section on cognitive data analysis, and Figure 5).

Next, to explore change in cognition over time and its relation to neural indices, we combined the LC–verbal learning models of both time points (T1 and T2) in a latent change score model^55, 58^ (see Supplementary Figure 10). As only cognitive performance was assessed at both time points, we calculated a univariate (multiple indicator) latent change score model. In this, we estimated the change in latent intercept and slope factors on a second order latent level, expressed by Δ-intercept and Δ-slope factors. Since the same word list was tested in all learning trials in the verbal learning memory task and an identical list was used at T1 and T2, we did not include residual covariances of error terms across time points or correlated errors (cf. ^58^). This model allows conclusions about the average rate of change in cognition for each age group (mean Δ-*) as well as about the within-group variance in change (variance Δ-*;^58^). The association between the cognitive latent change scores and the latent neural variables was assessed by means of freely estimated covariances. As before, model fit for all described models was determined using a *χ*^2^-test in combination with two additional fit indices (RMSEA, CFI; see Supplementary Results 2.3.2).

### Statistical parameters

All reported results are based on two-sided statistical tests. We used an alpha level of .05 for all statistical tests. Statistical results with *p*-values between 0.05 and 0.1 are described as statistical trend. When indicated, tests were adjusted for multiple comparisons. The applied statistical tests did not include covariates, unless noted differently. Data distribution was assumed to be normal (cf. Supplementary Figure 12) but this was not formally tested.

### Code availability

The custom code used for these analyses is available from the corresponding authors upon request.

### Data availability

The data that our results are based on are available from the <u>BASE-II steering committee</u> upon approved research proposal. For inquiries please contact Dr. Katrin Schaar, BASE-II project coordinator. To facilitate comparability of study results, we freely share the established LC probability map with the neuroscientific community (https://www.mpib-berlin.mpg.de/LC-Map).

## Acknowledgements

This article uses data from the Berlin Aging Study II (BASE-II), which was supported by the German Federal Ministry of Education and Research (Bundesministerium für Bildung und Forschung, BMBF) under grant numbers #16SV5536 K, #16SV5537, #16SV5538, and #16SV5837, #01UW070 and #01UW0808. Additional contributions (e.g., financial, equipment, logistics, personnel) are made from each of the other participating sites, i.e., the Max Planck Institute for Human Development (MPIB), Max Planck Institute for Molecular Genetics (MPIMG), Charite-Universiätsmedizin, University Medicine, German Institute for Economic Research (DIW), Humboldt-Universität zu Berlin, all located in Berlin, Germany, and University of Lübeck, and University of Tübingen, Germany. For further information about the BASE-II project, see https://www.base2.mpg.de/en

MW-B received support from the German Research Foundation (DFG, WE 4269/5-1) and the Jacobs Foundation (Early Career Research Fellowship 2017–2019).

MJD is a fellow of the International Max Planck Research School on the Life Course (LIFE; http://www.imprs-life.mpg.de/en). MJD is recipient of a stipend from the Sonnenfeld-Foundation (http://www.sonnenfeld-stiftung.de/en/). MM’s work was supported by an Alexander von Humboldt fellowship and by National Institutes of Health grant R01AG025340. The funders had no role in study design, data collection and analysis, decision to publish, or preparation of the manuscript.

We thank Andrew R. Bender, Matthew Betts, and Myriam C. Sander for valuable discussions and assistance. We are grateful to Shelby Bachman and Dilara Zorbek, who performed the manual tracing of the locus coeruleus, as well as Ylva Köhncke and Yana Fandakova for statistical advice, and Michael Krause for help with cluster computing.

## Author contributions

UL and SK designed the broader BASE-II study; MJD, MWB, MM, SK, and NCB designed the additional LC component; SD performed the experiments; MJD and MWB analyzed the data; MJD and MWB wrote the manuscript; UL, SK, MM, SD, and NCB gave conceptual advice. All authors revised the manuscript.

