## Supplementary Information for "Higher rostral locus coeruleus integrity is associated with better memory performance in older adults"

2.3.2 Associations between latent locus coeruleus integrity scores and change in memory performance 27

### 1 Supplementary tables and figures

**Supplementary Table 1.** Summary of sample descriptives for younger (YA) and older adults (OA).

| | YA (n = 66; 22 female) | | | | OA (n = 228; 82 female) | | | |
|---|---|---|---|---|---|---|---|---|
| | Mean | *SD* | Min | Max | Mean | *SD* | Min | Max |
| Age (at time point 2) | 32.5 | 3.53 | 25.41 | 39.84 | 72.29 | 4.11 | 62.53 | 83.16 |
| BMI | 23.02 | 3.78 | 11.59 | 31.86 | 26.75 | 3.49 | 19.13 | 37.92 |
| Education | 14.31 | 2.55 | 10 | 18 | 14.12 | 2.95 | 7 | 18 |
| DSST | 62.14 | 10.93 | 40 | 89.5 | 44.05 | 9.56 | 16.5 | 72 |
| MMSE | - | - | - | - | 28.57 | 1.3 | 22 | 30 |

*Note*. SD = Standard deviation; BMI = Body Mass Index; DDST = Digit Symbol Substitution Test; MMSE = Mini Mental State Examination. Age and education is expressed in years.

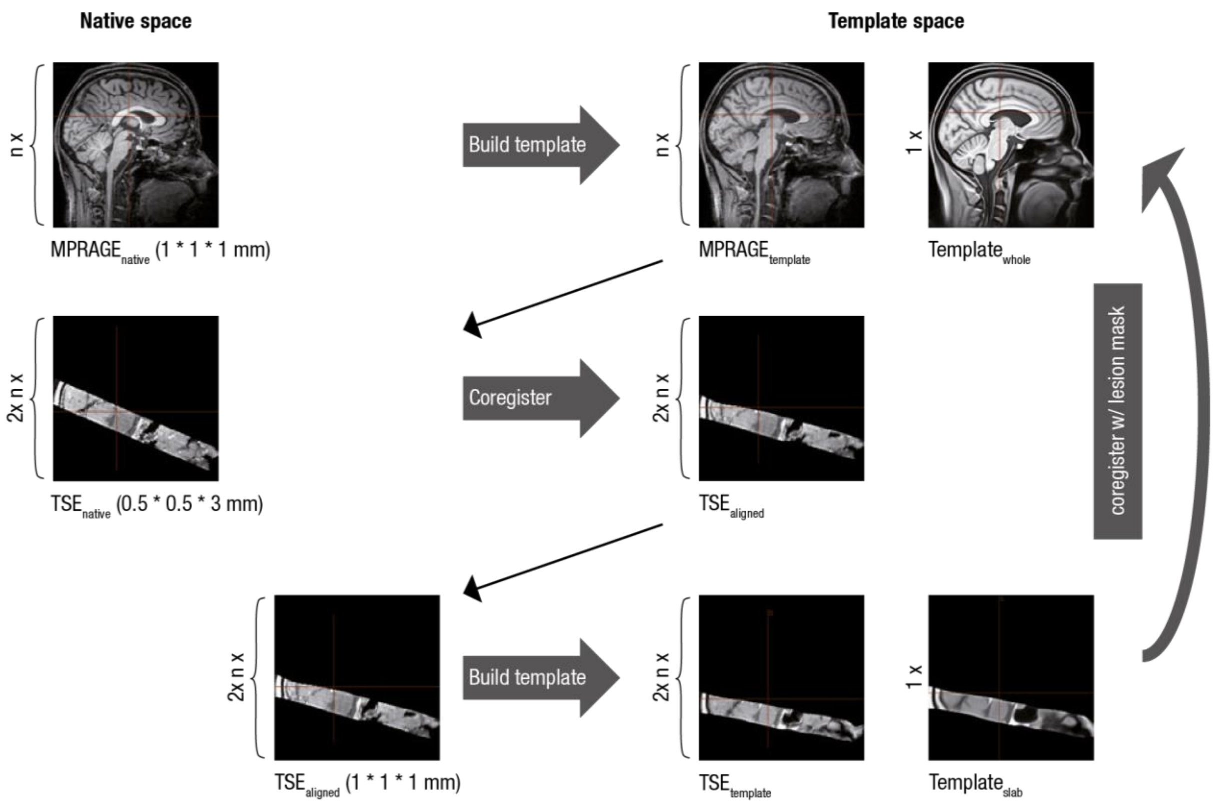

**Supplementary Figure 1.** Overview of coregistration and template building steps used to generate a brainstem template. Step 1 (top row): Individual whole brain scans ($MPRAGE_{native}$) are aligned within a template building algorithm and used to generate a group whole brain template ($Template_{whole}$). Step 2 (middle row): Within subjects we coregister native space brainstem scans to whole brain scans ($MPRAGE_{template}$) to align scans across subjects while maintaining high coregistration accuracy. Step 3 (bottom row): Aligned brainstem scans ($TSE_{aligned}$) are used to generate a group brainstem template ($Template_{slab}$) with increased signal-to-noise ratio. Brainstem and whole brain templates are coregistered to facilitate transformation to standard space.

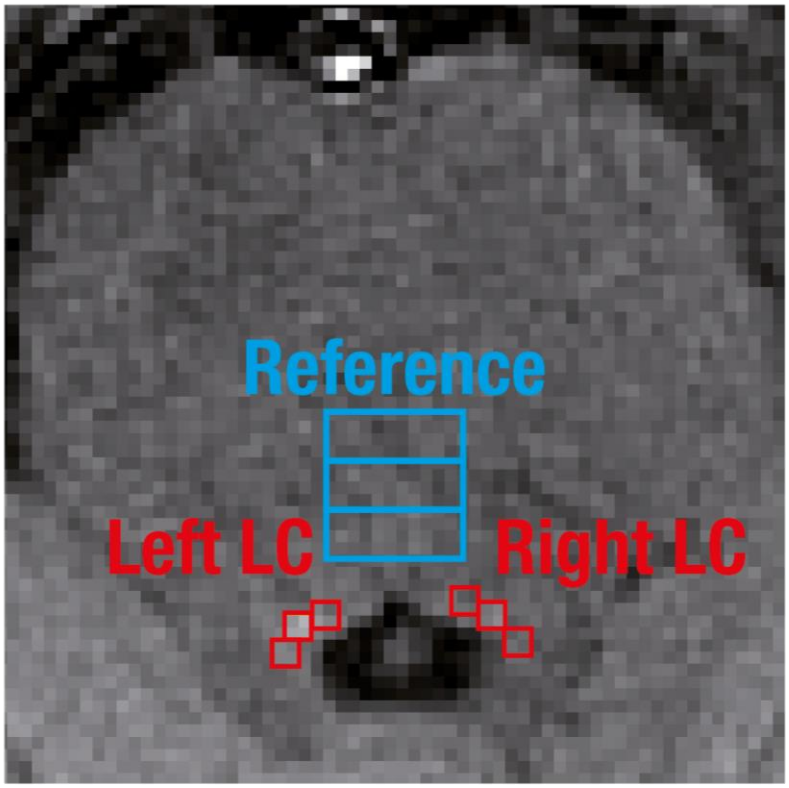

**Supplementary Figure 2.** Depiction of the manual locus coeruleus (LC; red) and reference (blue) intensity assessment. Two independent, blinded raters place rectangular regions of interest (ROI) to capture LC-neuromelanin-related hyperintensities next to the fourth ventricle. Reference intensity is assessed in the dorsal pontine tegmentum between the LC ROI (see Methods, section on comparison to manually assessed locus coeruleus intensity, for a full description of the rating procedure).

#### 2 Supplementary results

##### 2.1 Cognitive results

###### 2.1.1 Adequacy of the verbal learning and memory models

We captured the non-linear performance trajectories of younger and older adults' performance in the Rey Auditory Verbal Learning and Memory Task[1] (RAVLT) using a structural equation model. The proposed multiple-group quadratic growth curve model fits the data well both for time point 1 (T1) ($\chi^2(46) = 35.535$, RMSEA = 0.0, CFI = 1.01) and time point 2 (T2) ($\chi^2(46) = 81.764$, RMSEA = 0.052, CFI = 0.965 see Supplementary Figure 3). Further, the quadratic growth model showed considerably better fit to the data than competing alternative models (i.e., intercept only, linear, hyperbolic[2], logarithmic[3], and exponential slope models) as confirmed by likelihood ratio difference tests (for nested models) and comparisons of the Akaike Information Criterion (AIC; for non-nested models; see Supplementary Table 2). The quadratic growth curve models for T1 and T2 demonstrated strict factorial invariance (comparison of models with and without variant manifest errors (eV) using likelihood ratio tests: all $\Delta\chi^2(\Delta df = 1) \leq 2.727$; all $p \geq 0.099$).

**Supplementary Table 2.** Comparison of competing alternative growth models for time point 2

| Model | $\chi^2$ | $df$ | $\Delta\chi^2$ | $\Delta df$ | $p$ | RMSEA | CFI | AIC | RL |
|---|---|---|---|---|---|---|---|---|---|
| Icept | 1558.624 | 60 | 1476.861 | 14 | <0.001 | 0.292 | 0.0 | 7407.831 | -- |
| Icept + Lin | 405.838 | 54 | 324.075 | 8 | <0.001 | 0.149 | 0.657 | 6267.045 | -- |
| Icept+Lin+Exp | 140.126 | 46 | -- | -- | -- | 0.084 | 0.908 | 6017.333 | <0.001 |
| Hyperbolic | 90.052 | 46 | -- | -- | -- | 0.057 | 0.957 | 5967.259 | 0.016 |
| Icept+Lin+Log | 82.168 | 46 | -- | -- | -- | 0.052 | 0.965 | 5959.375 | 0.817 |
| Icept+Lin+Quad | 81.764 | 46 | -- | -- | -- | 0.052 | 0.965 | 5958.97 | -- |

*Note:* Icept = Intercept; Lin = Linear slope, Quad = Quadratic slope; Exp = Exponential slope; Log = Logarithmic slope; RMSEA = root mean square error of approximation; CFI =comparative fit index; AIC = Akaike Information Criterion; RL = Relative likelihood. Likelihood-ratio difference tests and relative likelihood tests compare the respective model to the Icept+Lin+Quad model.

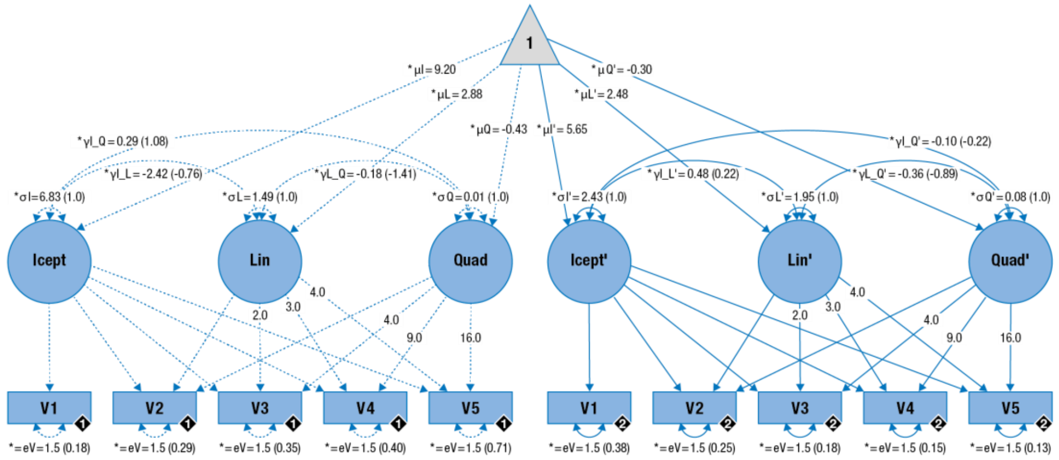

**Supplementary Figure 3.** Pictorial rendition of the structural equation model that approximates participants' learning curves with a quadratic function consisting of an initial memory performance level (intercept) and a gain over learning trials (linear and quadratic slope). Cognitive manifest variables represent the iteratively assessed memory performance in the verbal learning and memory task (V1–5). Black diamonds on manifest variables indicate the age group (younger adults = 1, n = 66, broken lines; older adults = 2, n = 228, solid lines). (Co)Variances ($\gamma$, $\sigma$) in brackets indicate standardized estimates. Loadings that are freely estimated (*) but constrained to be equal across age groups (=) are indicated by both asterisk and equal signs (*=). Icept = Intercept; Lin/Quad= linear / quadratic slope, respectively. Rectangles and circles indicate manifest and latent variables, respectively. The constant is depicted by a triangle.

##### 2.1.2 Verbal learning and memory within younger and older adults

For both time points and age groups, we observed reliable average intercept and slope factors (likelihood ratio tests: all $\Delta\chi^2(\Delta df = 1) \geq 24.39$, all $p < 0.001$; see Supplementary Table 3). Further, there were consistent interindividual differences in initial recall (intercept) and learning (quadratic and linear slope) parameters. This applied to both age groups at both time points as indicated by significant variances of the latent factors (likelihood ratio tests: all $\Delta\chi^2(\Delta df = 1) \geq 11.97$, all $p < 0.001$; see Supplementary Table 3). Only the variance of the quadratic slope factor failed to reach significance in younger adults at T2 (likelihood ratio tests: $\Delta\chi^2(\Delta df = 1) = 0.276$, $p = 0.599$, estimate (est) = 0.011, [95% confidence interval (CI): -0.031, 0.053]).

The covariances between initial recall and learning rate (e.g., starting out higher is related to faster performance increases) demonstrated a mixed pattern. While for T1, there was no statistically significant association between initial recall memory and learning rate (i.e., the covariance between intercept and quadratic/linear slope factors; likelihood ratio tests: all $\Delta\chi^2(\Delta df = 1) \leq 1.603$, all $p \geq 0.205$; see Supplementary Table 3), there was a reliable association at T2 for younger but not older adults (likelihood ratio tests; covariance intercept and quadratic slope in younger adults: $\Delta\chi^2(\Delta df = 1) = 6.656$, $p = 0.01$, $est_{younger\ adults} = 0.293$, [95% CI: 0.026, 0.561]; covariance intercept and quadratic slope in older adults: $\Delta\chi^2(df = 1) = 2.593$, $p = 0.107$, $est_{older\ adults} = -0.097$, [95% CI: -0.210, 0.017]. Covariance intercept and linear slope in younger adults: $\Delta\chi^2(\Delta df = 1) = 19.076$, $p < 0.001$, $est_{younger\ adults} = -2.420$, [95% CI: -3.954, -0.885]; covariance intercept and linear slope in older adults: $\Delta\chi^2(\Delta df = 1) = 3.049$, $p = 0.081$, $est_{older\ adults} = 0.482$, [95% CI: -0.034, 0.998]). Further, a steeper linear increase in memory performance was reliably associated with a more negative acceleration of growth at T1 for younger and older adults (i.e., the covariance between linear and quadratic slope factors; likelihood ratio tests: all $\Delta\chi^2(\Delta df = 1) \geq 20.453$, all $p \leq 0.001$; see Supplementary Table 3). At T2, we observed a marginally significant statistical effect (likelihood ratio test for younger adults: $\Delta\chi^2(\Delta df = 1) = 3.788$, $p = 0.052$ , $est_{younger\ adults} = -0.178$, [95% CI: -0.395, 0.038]) and a reliable association between slope factors for older adults (likelihood ratio test: $\Delta\chi^2(\Delta df = 1) = 30.484$, $p < 0.001$, $est_{older\ adults} = -0.358$, [95% CI: -0.516, -0.200]).

**Supplementary Table 3**. Overview of verbal learning and memory performance within younger and older adults for time point 1 and 2

| | Parameter | TP | Group | $\Delta\chi^2$ | $\Delta df$ | $p$ | est | CI lower | CI upper |
|---|---|---|---|---|---|---|---|---|---|
| **μ** | Icept | 1 | YA | 151.671 | 1 | <0.001 | 8.421 | 7.758 | 9.084 |
| **μ** | Lin | 1 | YA | 54.389 | 1 | <0.001 | 2.566 | 2.023 | 3.109 |
| **μ** | Quad | 1 | YA | 24.390 | 1 | <0.001 | -0.327 | -0.445 | -0.209 |
| **σ** | Icept | 1 | YA | 113.001 | 1 | <0.001 | 5.857 | 3.317 | 8.396 |
| **σ** | Lin | 1 | YA | 27.966 | 1 | <0.001 | 2.872 | 1.152 | 4.591 |
| **σ** | Quad | 1 | YA | 15.651 | 1 | <0.001 | 0.113 | 0.032 | 0.194 |
| γ | Icept–Lin | 1 | YA | 0.01 | 1 | 0.921 | -0.076 | -1.589 | 1.437 |
| γ | Lin–Quad | 1 | YA | 20.453 | 1 | <0.001 | -0.555 | -0.922 | -0.189 |
| γ | Icept–Quad | 1 | YA | 1.603 | 1 | 0.205 | -0.214 | -0.533 | 0.106 |
| **μ** | Icept | 1 | OA | 494.03 | 1 | <0.001 | 5.571 | 5.314 | 5.828 |
| **μ** | Lin | 1 | OA | 137.486 | 1 | <0.001 | 2.087 | 1.790 | 2.384 |
| **μ** | Quad | 1 | OA | 48.959 | 1 | <0.001 | -0.235 | -0.297 | -0.173 |
| **σ** | Icept | 1 | OA | 84.741 | 1 | <0.001 | 2.387 | 1.655 | 3.118 |
| **σ** | Lin | 1 | OA | 76.238 | 1 | <0.001 | 3.095 | 2.115 | 4.076 |
| **σ** | Quad | 1 | OA | 37.51 | 1 | <0.001 | 0.109 | 0.065 | 0.153 |
| γ | Icept–Lin | 1 | OA | 0.1 | 1 | 0.752 | 0.101 | -0.518 | 0.720 |
| γ | Lin–Quad | 1 | OA | 46.008 | 1 | <0.001 | -0.524 | -0.724 | -0.325 |
| γ | Icept–Quad | 1 | OA | 0.088 | 1 | 0.767 | -0.020 | -0.148 | 0.109 |
| **μ** | Icept | 2 | YA | 160.461 | 1 | <0.001 | 9.197 | 8.508 | 9.886 |
| **μ** | Lin | 2 | YA | 82.158 | 1 | <0.001 | 2.880 | 2.438 | 3.322 |
| **μ** | Quad | 2 | YA | 62.602 | 1 | <0.001 | -0.432 | -0.515 | -0.349 |
| **σ** | Icept | 2 | YA | 153.377 | 1 | <0.001 | 6.825 | 4.039 | 9.612 |
| **σ** | Lin | 2 | YA | 11.97 | 1 | <0.001 | 1.491 | 0.326 | 2.655 |
| **σ** | Quad | 2 | YA | 0.276 | 1 | 0.599 | 0.011 | -0.031 | 0.053 |
| γ | Icept–Lin | 2 | YA | 19.076 | 1 | <0.001 | -2.420 | -3.954 | -0.885 |
| γ | Lin–Quad | 2 | YA | 3.788 | 1 | 0.052 | -0.178 | -0.395 | 0.038 |
| γ | Icept–Quad | 2 | YA | 6.656 | 1 | 0.010 | 0.293 | 0.026 | 0.561 |
| **μ** | Icept | 2 | OA | 512.977 | 1 | <0.001 | 5.647 | 5.395 | 5.899 |
| **μ** | Lin | 2 | OA | 218.765 | 1 | <0.001 | 2.477 | 2.224 | 2.730 |
| **μ** | Quad | 2 | OA | 87.829 | 1 | <0.001 | -0.299 | -0.355 | -0.242 |
| **σ** | Icept | 2 | OA | 99.634 | 1 | <0.001 | 2.429 | 1.722 | 3.135 |
| **σ** | Lin | 2 | OA | 45.819 | 1 | <0.001 | 1.945 | 1.214 | 2.677 |
| **σ** | Quad | 2 | OA | 29.039 | 1 | <0.001 | 0.083 | 0.046 | 0.120 |
| γ | Icept–Lin | 2 | OA | 3.049 | 1 | 0.081 | 0.482 | -0.034 | 0.998 |
| γ | Lin–Quad | 2 | OA | 30.484 | 1 | <0.001 | -0.358 | -0.516 | -0.200 |
| γ | Icept–Quad | 2 | OA | 2.593 | 1 | 0.107 | -0.097 | -0.210 | 0.017 |

Note: Icept = Intercept; Lin = Linear slope, Quad = Quadratic slope; μ = Mean; σ = Variance; γ = Covariance; TP = Time Point; YA = Younger adults (n = 66); OA = Older adults (n = 228); *df* = degrees of freedom; est = parameter estimate; CI = 95 % confidence interval; All statistical comparisons are based on likelihood ratio tests. Also see Supplementary Figure 3 for a model depiction.

##### 2.1.3 Age differences in verbal learning and memory

At T1, likelihood ratio difference tests revealed significant age differences in the intercept factors (likelihood ratio test: $\Delta\chi^2(\Delta df = 1) = 45.391$, $p < 0.001$, $est_{younger\ adults} = 8.421$, [95% CI: 7.758, 9.084], $est_{older\ adults} = 5.571$, [95% CI: 5.314, 5.828]), pointing to higher initial recall performance in younger adults. However, age differences in the slope factors failed to reach significance (likelihood ratio tests; linear: $\Delta\chi2(\Delta df = 1) = 2.274$, $p = 0.132$, $est_{younger\ adults} = 2.566$, [95% CI: 2.023, 3.109], $est_{older\ adults} = 2.087$, [95% CI: 1.790, 2.384]; quadratic: $\Delta\chi^2(\Delta df = 1) = 1.819$, $p = 0.177$, $est_{younger\ adults} = -0.327$, [95% CI: -0.445, -0.209], $est_{older\ adults} = -0.235$, [95% CI: -0.297, -0.173]). We observed no statistically reliable age differences in the covariances between the latent intercept and slope factors (likelihood ratio tests: all $\Delta\chi^2(df = 1) \leq 1.191$, all $p \geq 0.275$; see Supplementary Table 4).

For T2, likelihood ratio difference tests also indicated age differences in the intercept factors (likelihood ratio tests: $\Delta\chi^2(\Delta df = 1) = 59.533$, $p < 0.001$, $est_{younger\ adults} = 9.197$, [95% CI: 8.508, 9.886], $est_{older\ adults} = 5.647$, [95% CI: 5.395, 5.899]). Again, we observed no significant difference in the linear slope factors between groups (likelihood ratio test: $\Delta\chi^2(\Delta df = 1) = 2.381$, $p = 0.123$, $est_{younger\ adults} = 2.880$, [95% CI: 2.438, 3.322], $est_{older\ adults} = 2.477$, [95% CI: 2.224, 2.730]), however, the difference in the quadratic slope term turned out to be significant (likelihood ratio test: $\Delta\chi^2(\Delta df = 1) = 6.579$, $p = 0.01$, $est_{younger\ adults} = -0.432$, [95% CI: -0.515, -0.349], $est_{older\ adults} = -0.299$, [95% CI: -0.355, -0.242]). This finding points to age differences in the negative acceleration of the learning curves. Note however, that ceiling effects may play a role here (see main text, Figure 1). The associations between initial recall and learning factors differed reliably among younger and older adults (likelihood ratio tests; covariance intercept and quadratic slope: $\Delta\chi^2(\Delta df = 1) = 9.454$, $p = 0.002$, $est_{younger\ adults} = 0.293$, [95% CI: 0.026, 0.561],

$est_{older\ adults}$ = -0.097, [95% CI: -0.210, 0.017]. Covariance intercept and linear slope: $\Delta\chi^2(\Delta df = 1)$ = 21.839, $p < 0.001$, $est_{younger\ adults}$ = -2.420, [95% CI: -3.954, -0.885], $est_{older\ adults}$ = 0.482, [95% CI: -0.034, 0.998]). However, we observed no reliable age differences in the relation of the slope factors (likelihood ratio test: $\Delta\chi^2(\Delta df = 1)$ = 1.571, $p = 0.21$, $est_{younger\ adults}$ = -0.178, [95% CI: -0.395, 0.038], $est_{older\ adults}$ = -0.358, [95% CI: -0.516, -0.200]).

In sum, a quadratic growth model adequately described the observed learning curves of younger and older adults' performance in the RAVLT for both time points. Within groups, we observed reliable average initial recall (intercept) and learning (slope) factors. Individuals in both age groups differed in their initial recall level and the rate of learning. Age differences were mainly observed in initial recall performance.

**Supplementary Table 4**. Overview of age differences in verbal learning and memory performance for time point 1 and 2

| | Parameter | TP | Comparison | $\Delta\chi^2$ | $\Delta df$ | *p* |
|---|---|---|---|---|---|---|
| **μ** | Icept | 1 | YA vs OA | 45.391 | 1 | <0.001 |
| **μ** | Lin | 1 | YA vs OA | 2.274 | 1 | 0.132 |
| **μ** | Quad | 1 | YA vs OA | 1.819 | 1 | 0.177 |
| **σ** | Icept | 1 | YA vs OA | 11.485 | 1 | <0.001 |
| **σ** | Lin | 1 | YA vs OA | 0.049 | 1 | 0.825 |
| **σ** | Quad | 1 | YA vs OA | 0.009 | 1 | 0.923 |
| **μ** | Icept | 2 | YA vs OA | 59.533 | 1 | <0.001 |
| **μ** | Lin | 2 | YA vs OA | 2.381 | 1 | 0.525 |
| **μ** | Quad | 2 | YA vs OA | 6.579 | 1 | 0.021 |
| **σ** | Icept | 2 | YA vs OA | 17.437 | 1 | <0.001 |
| **σ** | Lin | 2 | YA vs OA | 0.403 | 1 | 0.21 |
| **σ** | Quad | 2 | YA vs OA | 5.33 | 1 | 0.002 |

Note: Icept = Intercept; Lin = Linear slope, Quad = Quadratic slope; μ = Mean; σ = Variance; TP = Time Point; YA = Younger adults (n = 66); OA = Older adults (n = 228); *df* = degrees of freedom; All statistical comparisons are based on likelihood ratio tests. For parameter estimates and confidence intervals, please see Supplementary Table 3.

##### 2.1.4 Adequacy of the general episodic memory model

To obtain a single measure for general episodic memory performance, i.e., independent of the specific task used, we made use of the comprehensive cognitive battery available for this data set (see Methods, section on cognitive data assessment). In particular, using an established episodic memory factor structure[4] we repeated our analyses (T2 only), this time integrating performance across multiple memory tasks while explicitly excluding information from the RAVLT (see Supplementary Figure 4). The model adequately fit the observed data ($\chi^2(14) = 1.991$, RMSEA = 0.0, CFI = 1.312). However, the model only demonstrated weak factorial invariance (i.e., it required variant manifest intercepts across groups) and thus precluded an interpretation of age group differences in the means of latent factors[5,6] (comparison of models with and without variant manifest intercepts (e.g., λFPT), likelihood ratio test: $\Delta\chi^2(\Delta df = 2) = 19.799$; $p < 0.001$). For both age groups, we observed reliable average episodic memory factors (likelihood ratio tests: all $\Delta\chi^2(\Delta df = 1) \geq 134.486$; all $p < 0.001$, see Supplementary Table 5) as well as interindividual differences therein (likelihood ratio tests: all $\Delta\chi^2(\Delta df = 1) \geq 29.606$; all $p < 0.001$, see Supplementary Table 5). In sum, the general episodic memory model adequately described the observed data. Within groups, we detected reliable latent episodic memory factors and individual differences therein.

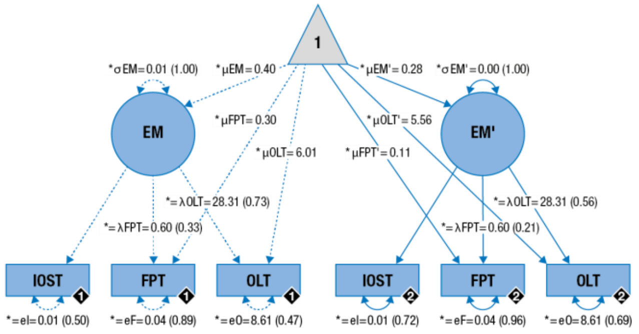

**Supplementary Figure 4.** Graphical depiction of the structural equation model that captures participants' general episodic memory performance in a single latent factor. Cognitive manifest variables represent memory performance in the indoor/outdoor scene encoding task (IOST), face–profession task (FPT), and object–location task (OLT). Black diamonds on manifest variables indicate the age group (younger adults = 1, n = 66, broken lines; older adults = 2, n = 228, solid lines). (Co)Variances (*γ, σ*) and loadings (*λ*) in brackets indicate standardized estimates. Loadings that are freely estimated (*) but constrained to be equal across age groups (=) are indicated by both asterisk and equal signs (*=). Rectangles and circles indicate manifest and latent variables, respectively. The constant is depicted by a triangle.

**Supplementary Table 5.** Overview of general episodic memory performance within younger and older adults for time point 2

| | Parameter | Group | $\Delta\chi^2$ | $\Delta df$ | *p* | est | CI lower | CI upper |
|---|---|---|---|---|---|---|---|---|
| **μ** | EM | YA | 134.486 | 1 | <0.001 | 0.398 | 0.361 | 0.436 |
| **σ** | EM | YA | 47.749 | 1 | <0.001 | 0.012 | 0.003 | 0.022 |
| **μ** | EM | OA | 442.803 | 1 | <0.001 | 0.281 | 0.264 | 0.298 |
| **σ** | EM | OA | 20.546 | 1 | <0.001 | 0.005 | 0.001 | 0.008 |

Note: EM = general episodic memory; μ = Mean; σ = Variance; YA = Younger adults (n = 66); OA = Older adults (n = 228); *df* = degrees of freedom; est = parameter estimate; CI = 95 % confidence interval; All statistical comparisons are based on likelihood ratio tests. Age group comparisons are omitted since the model demonstrated only weak factorial invariance.

#### 2.2 Magnetic resonance imaging results

##### 2.2.1 Validity of automatically assessed locus coeruleus intensity ratios

Before the investigation of age differences in automatically assessed locus coeruleus (LC) intensity and their relation to learning and memory performance, we sought to establish the validity of the proposed approach. Unless otherwise stated, all following reports concern LC measures automatically assessed in template space ($TSE_{template}$; see Methods, section on coregistration and standardization of magnetic resonance imaging data).

###### *2.2.1.1 Comparison to previously published locus coeruleus masks*

We semi-automatically extracted individual peak intensities across the rostrocaudal extent of the LC. Peak intensity coordinates were converted to a probability map and warped into 0.5 mm iso-voxel standard space where we calculated the overlap with previously published LC maps[7,8]. The thresholded probability map contains 175 voxels equivalent to a volume of 26 $mm^3$. It includes two mostly symmetrical cylindrical structures corresponding to left and right LC that shift somewhat more laterally and posteriorly, following the wall of the fourth ventricle, along the rostrocaudal axis. The voxels of the proposed map overlap largely with the map published by Betts and colleagues[8] (57.71%) as well as the one by Keren and others[7] (69.14%; see main text, Figure 3 d; we used Keren's 1 SD LC map for comparisons). About half of the map lies within the intersection of all three maps (40%; compared to 9.41% and 8.53% for Keren et al.[7] and Betts et al.[8], respectively). Thus, taken together, 86.86% of the probability map is in line with previously published maps. However, our probability map extends less caudally compared to the other two, which explains why proportionally fewer segments of their maps fall within its boundaries (12.30% and 16.26% of Betts et al.[8] and Keren et al.[7], respectively). The two reference maps themselves overlap moderately (22.78% of Betts et al. in Keren et al., and 25.13% for the inverse[7,8]).

###### 2.2.1.2 Comparison to manually assessed locus coeruleus intensity

Two independent, blinded raters (research assistants) manually assessed LC intensity on all axial slices that showed elevated signal in anatomically plausible LC locations. Manually assessed peak intensities showed high accordance between raters as demonstrated by means of intraclass correlation coefficients (ICC; left LC intensity: $F(289, 289) = 79.579$, $p < 0.001$, ICC = 0.987, [95 % CI: 0.984, 0.990]; right LC intensity: $F(289, 289) = 79.629$, $p < 0.001$, ICC = 0.987, [95 % CI: 0.984, 0.990]), so we collapsed intensity values across raters. On average, elevated signal intensity was detected on 3.028 slices ($SD = 0.122$, range = 3–4). Manually and automatically assessed LC intensities were closely related (intraclass correlations: left LC: $F(289, 289) = 28.615$, $p < 0.001$, ICC = 0.933, [95 % CI: 0.585, 0.975]; right LC: $F(289, 289) = 18.314$, $p < 0.001$, ICC = 0.896, [95 % CI: 0.430, 0.961]; mean over left and right LC: $F(289, 289) = 38.283$, $p < 0.001$, ICC = 0.931, [95 % CI: 0.263, 0.979]) highlighting the validity of the proposed method (though interindividual differences in overall MRI brightness may have inflated estimates).

###### 2.2.1.3 Reproducibility of semi-automatically assessed locus coeruleus intensity ratios

In order to judge the temporal stability of the semi-automatic method, we repeatedly over the course of several weeks scanned a small number of younger adults that did not participate in the main study. We applied the same coregistration and template building steps as in the main study (see Methods, section on coregistration and standardization of magnetic resonance imaging data). After alignment with the study brainstem template ($Template_{slab}$), we used the established search spaces to extract intensity ratios. Across six measurements, we observed high reproducibility of automatically assessed peak LC ratios (intraclass correlation; mean over left and right hemisphere: $F(2, 10) = 8.052$, $p = 0.008$, ICC = 0.873, [95 % CI: 0.402, 0.997]) compared to manual assessments[9]. In addition, these results demonstrate the applicability of the semi-automatic procedure for small samples.

###### *2.2.1.4 Comparison to automatically assessed locus coeruleus intensity in native space*

We warped the LC and reference search spaces to individual subject's coordinates (i.e., native space) and extracted intensity values for all participants. Values automatically assessed in template and native space demonstrated high accordance (intraclass correlations; mean over left and right LC: $F(293, 293) = 65.261$, $p < 0.001$, ICC = 0.906, [95 % CI: 0.0, 0.977]; left LC: $F(293, 293) = 51.813$, $p < 0.001$, ICC = 0.909, [95 % CI: 0.0, 0.976]; right LC: $F(293, 293) = 39.022$, $p < 0.001$, ICC = 0.896, [95 % CI: 0.0, 0.972]) indicating the stability of the proposed method. However, we note that the obtained ICCs show wide confidence intervals, potentially resulting from changes in absolute LC intensity by the entailed warping and registration steps. We thus repeated the analyses, now exploring the consistency in intensity values across measures (i.e., ICC; two-way mixed model with consistency between template and native space values; mean over left and right LC: $F(293, 293) = 65.261$, $p < 0.001$, ICC = 0.985, [95 % CI: 0.981, 0.988]; left LC: $F(293, 293) = 51.813$, $p < 0.001$, ICC = 0.981, [95 % CI: 0.976, 0.985]; right LC: $F(293, 293) = 39.022$, $p < 0.001$, ICC = 0.974, [95 % CI: 0.968, 0.980]). These follow-up analyses suggest that while transformation between native and template space may change absolute LC intensity, the rank order of values within the sample is stable.

##### 2.2.2 Age differences in the spatial distribution of locus coeruleus intensity ratios

A non-parametric cluster-based permutation test[10] (see Methods, section on analysis of age differences in the spatial distribution of locus coeruleus ratios) revealed spatially heterogeneous age differences in LC ratios along the rostrocaudal extent of the nucleus (see Supplementary Figure 5a). Relative to younger adults, older adults showed a cluster of elevated intensity spanning caudal slices (left LC: 59th–100th LC percentile; cluster permutation test: $p_{corr} < 0.001$, [95 % CI: <0.001, <0.001]; right LC: 65th–94th LC percentile; cluster permutation test: $p_{corr} = 0.001$, [95 % CI: 0.001, 0.001]). In contrast, older adults demonstrated a tendency towards reduced intensity values in rostral LC segments that, however, did not reach significance in the unilateral analyses (left LC: 29th–35th LC percentile; cluster permutation test: $p_{corr} = 0.139$, [95 % CI: 0.137, 0.140]; right LC: 29th–35th LC percentile; cluster permutation test: $p_{corr} = 0.107$, [95 % CI: 0.105, 0.108]). In general, these hemisphere-specific analyses replicate the bilateral findings reported in the main text and indicate spatially confined age differences in LC intensity[11].

Previous studies reported lateralized differences in LC ratios (left > right; cf.[8,9]) that we replicate here by means of Wilcoxon signed rank tests. In particular, we computed tests (collapsing over slices) across age groups, and then for each group separately (Wilcoxon signed rank tests; younger and older adults: $W(294) = 9397$, $Z = -8.421$, $p < 0.001$, $r = -0.4911$; [95 % CI: -0.573, -0.399][12]; younger adults: $W(66) = 628$, $Z = -3.050$, $p = 0.002$, $r = -0.375$, [95 % CI: -0.566, -0.147]; older adults: $W(228) = 5143$, $Z = -7.933$, $p < 0.001$, $r = -0.525$, [95 % CI: -0.613, -0.424]).

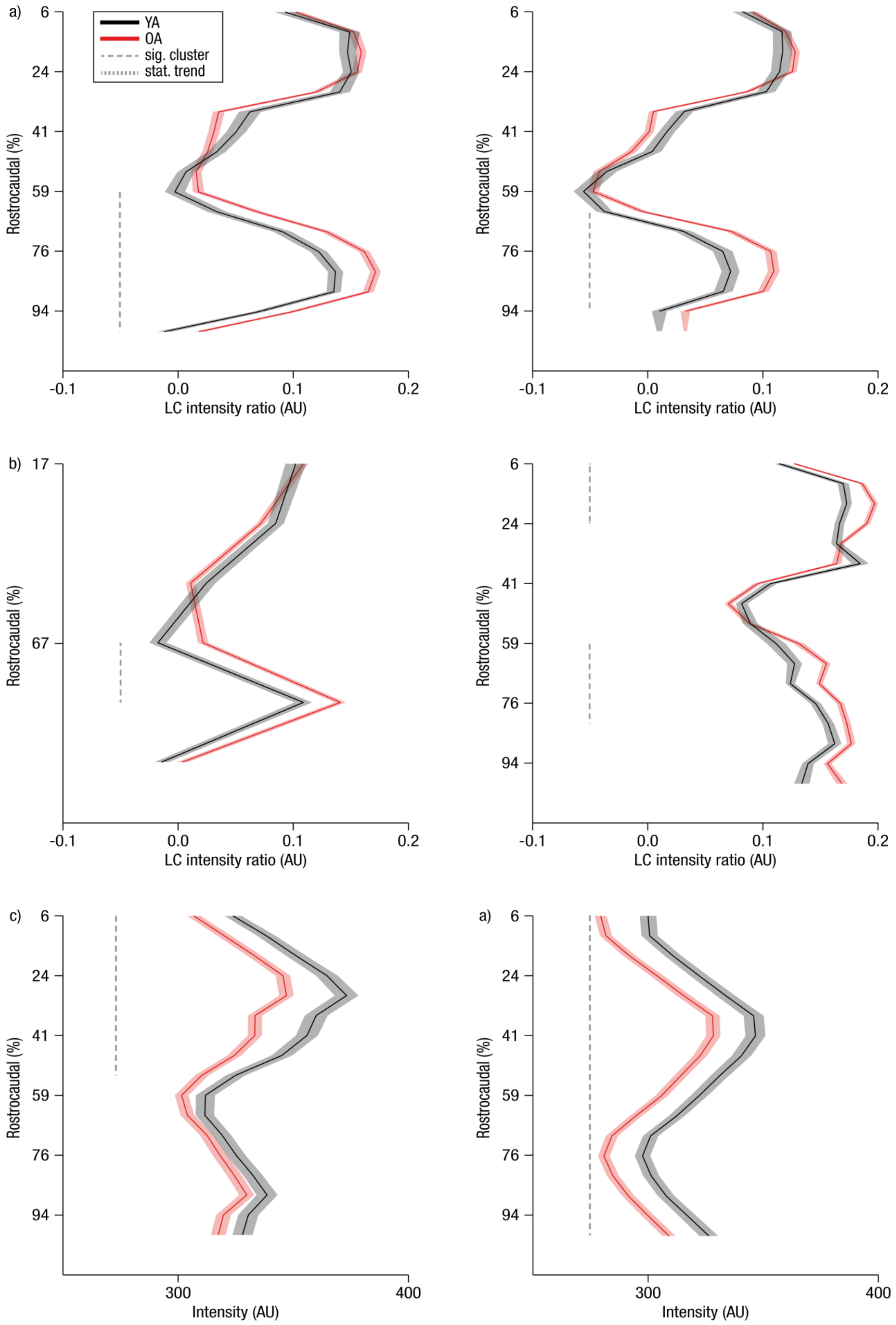
a)
b)
c)
a)
YA
OA
sig. cluster
stat. trend
Rostrocaudal (%)
6
24
41
59
76
94
17
67
-0.1
0.0
0.1
0.2
LC intensity ratio (AU)
300
400
Intensity (AU)

**Supplementary Figure 5**. Analyses of age differences in locus coeruleus (LC) intensity ratios (a–b) and raw intensity (c) across the rostrocaudal axis by means of cluster-based permutation tests. (a) Peak intensity ratios automatically assessed in template space were analyzed separately for the left and right hemisphere (left, right, respectively). (b) Comparisons based on peak native space (left plot) and mean template space intensity (right plot) LC measures. (c) Analysis of peak raw intensity values split for LC and reference (left, right, respectively). Shaded areas represent ± 1 standard error of the mean. Data of younger (n = 66; black lines) and older adults (n = 288; red lines) is presented.

#### 2.2.3 Estimation of latent locus coeruleus integrity scores

##### *2.2.3.1 Adequacy of the latent locus coeruleus integrity model*

We used a multiple-group single factor SEM to estimate overall LC integrity scores on a latent level while accounting for measurement error in the observed variables (see Supplementary Figure 6). For each age group, peak LC ratios (averaged across slices; see Methods, section on estimation of latent locus coeruleus integrity scores) of the left and right hemispheres loaded on a single LC factor. Before model estimation, LC ratios were scaled (multiplied × 100) to facilitate model estimation. In the model, error variances, manifest means, and factor loadings were constrained to be equal across age groups (i.e., strict factor invariance; comparison of models with and without variant manifest errors, means, and factor loadings; likelihood ratio test: $\Delta\chi^2(\Delta df = 3) = 1.791$; $p = 0.617$). The proposed model fit the data well ($\chi^2(7) = 1.791$, RMSEA = 0.000, CFI = 1.222).

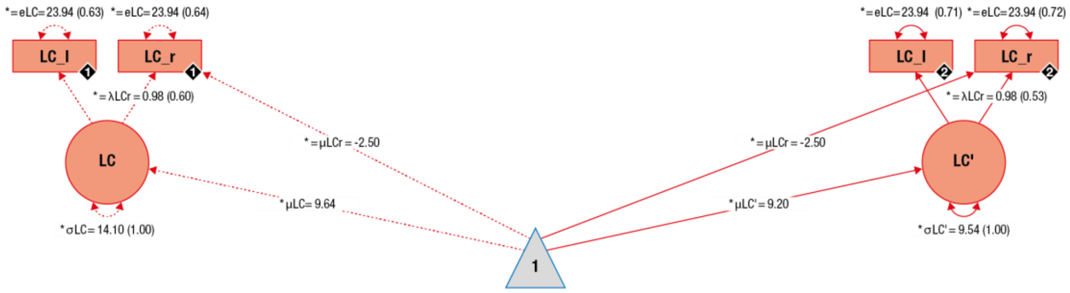

Supplementary Figure 6. Graphical depiction of the structural equation model that integrates over locus coeruleus intensity data to estimate a single locus coeruleus integrity factor (LC; red) on a latent level for younger and older adults. Neural manifest variables are the LC intensity ratios of each hemisphere (left = LC_l; right = LC_r). Black diamonds on manifest variables indicate the age group (younger adults = 1, n = 66, broken lines; older adults = 2, n = 228, solid lines). (Co)Variances (γ, σ) and loadings (λ) in brackets indicate standardized estimates. Loadings that are freely estimated (*) but constrained to be equal across age groups (=) are indicated by both asterisk and equal signs (*=). Rectangles and circles indicate manifest and latent variables, respectively. The constant is depicted by a triangle.

##### *2.2.3.2 Latent locus coeruleus integrity scores within younger and older adults*

We detected reliable average latent LC factors for both age groups (likelihood ratio tests: all $\Delta\chi^2(\Delta df = 1) \geq 90.454$, all $p < 0.001$, see Supplementary Table 6), and within each group, subjects showed significant interindividual differences in LC scores (likelihood ratio tests: all $\Delta\chi^2(\Delta df = 1) \geq 18.305$, all $p < 0.001$; see Supplementary Table 6).

Supplementary Table 6. Overview of locus coeruleus integrity scores within younger and older adults

| | Parameter | Group | $\Delta\chi^2$ | $\Delta df$ | $p$ | est | CI lower | CI upper |
|---|---|---|---|---|---|---|---|---|
| **μ** | LC | YA | 90.454 | 1 | <0.001 | 9.64048 | 8.342 | 10.939 |
| **σ** | LC | YA | 18.305 | 1 | <0.001 | 14.10496 | 3.957 | 24.253 |
| **μ** | LC | OA | 289.869 | 1 | <0.001 | 9.20499 | 8.484 | 9.926 |
| **σ** | LC | OA | 21.313 | 1 | <0.001 | 9.54461 | 3.787 | 15.302 |
| **μ** | LC | YA vs OA | 0.383 | 1 | 0.536 | -- | | |
| **σ** | LC | YA vs OA | 0.952 | 1 | 0.329 | -- | | |

Note: LC = locus coeruleus; μ = Mean; σ = Variance; YA = Younger adults (n = 66); OA = Older adults (n = 228); *df* = degrees of freedom; est = parameter estimate; CI = 95 % confidence interval; All statistical comparisons are based on likelihood ratio tests.

###### *2.2.3.3 Age differences in latent locus coeruleus integrity scores*

Age group differences in average LC scores did not reach statistical significance (likelihood ratio test: $\Delta\chi^2(\Delta df = 1) = 0.383$, $p = 0.536$, $est_{younger\ adults} = 9.640$, [95% CI: 8.342, 10.939], $est_{older\ adults} = 9.205$, [95% CI: 8.484, 9.926]) compatible with spatially confined age differences in LC integrity[7,8]. In sum, a multiple-group, single factor model adequately captures the interindividual differences in LC intensity ratios in the data. Within each age group, we detected reliable latent LC integrity factors as well as interindividual differences within them.

#### 2.3 Combined cognitive and magnetic resonance imaging results

##### 2.3.1 Associations between latent locus coeruleus integrity scores and memory performance

After establishing valid models for memory performance (see Supplementary Figures 3 and 4) and for LC integrity (see Supplementary Figure 6) in isolation, we sought to link the two modalities.

We first investigated the association between LC integrity and memory performance as assessed by the RAVLT. Next, to determine whether LC integrity is associated with episodic memory performance more generally, i.e., independent of the specific task used, we exploited the comprehensive cognitive battery available for this data set (see Methods, section on cognitive data assessment). Specifically, we repeated our analyses, using an established episodic memory factor structure[4] this time integrating performance across multiple memory tasks while explicitly excluding information of the RAVLT.

###### *2.3.1.1 Model adequacy*

We merged both modalities in unified neuro-cognitive models that demonstrated good fit for both of the cognitive measures used (i.e., RAVLT and general episodic memory; cf. Supplementary Table 7; see also main text, Figure 2, and Supplementary Figure 7).

Supplementary Table 7. Fit statistics for unified neural and cognitive models

| Model | | $\chi^2$ | *df* | RMSEA | CFI |
|---|---|---|---|---|---|
| LC–RAVLT | (Time point 1) | 47.41 | 87 | 0.0 | 1.037 |
| LC–RAVLT | (Time point 2) | 101.942 | 87 | 0.024 | 0.986 |
| LC–EM | (Time point 2) | 14.727 | 43 | 0.0 | 1.515 |

*Note:* LC = Locus Coeruleus; RAVLT = Rey Auditory Verbal Learning Task; EM = General Episodic Memory; df = Degrees of freedom; RMSEA = Root Mean Square Error of Approximation; CFI = Comparative Fit Index.

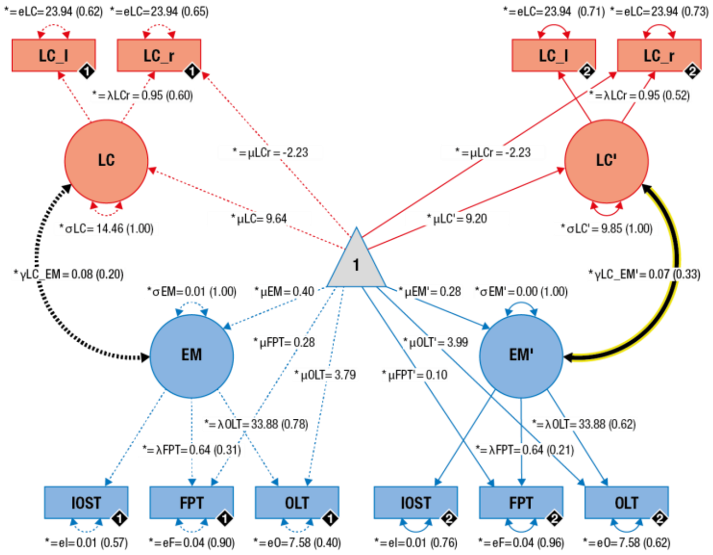

Supplementary Figure 7. Graphical depiction of the structural equation model that probes associations (thick black lines) between locus coeruleus integrity (LC; red) and general memory performance (blue) in younger and older adults on a latent level. Cognitive manifest variables represent the memory performance across three memory tasks (Indoor/Outdoor Scene Task (IOST); Face–Profession Task (FPT); Object–Location Task (OLT)). Neural manifest variables are the LC intensity ratios of each hemisphere (left = LC_l; right = LC_r). Black diamonds on manifest variables indicate the age group (younger adults = 1, n = 66, broken lines; older adults = 2, n = 228, solid lines). (Co)Variances ($\gamma$, $\sigma$) and loadings ($\lambda$) in brackets indicate standardized estimates. Loadings that are freely estimated (*) but constrained to be equal across age groups (=) are indicated by both asterisk and equal signs (*=). EM = General Episodic Memory. Rectangles and circles indicate manifest and latent variables, respectively. The constant is depicted by a triangle. The significant association between locus coeruleus integrity and memory performance (EM) in older adults is highlighted (yellow frame).

##### *2.3.1.2 Latent locus coeruleus integrity scores and memory performance within younger and older adults*

We observed a positive association between LC integrity and memory performance in older adults, irrespective of the cognitive measure used (i.e., RAVLT or general episodic memory) and time point analyzed (i.e., T1 or T2; see Supplementary Table 8).

In particular, at T1, older adults with higher LC integrity (assessed at T2) demonstrated higher initial recall performance (likelihood ratio test: $\Delta\chi^2(\Delta df = 1) = 3.993$, $p = 0.046$, est = 1.256, [95% CI: -0.020, 2.532], standardized est = 0.253 – please note that the estimation of confidence intervals focuses on a single parameter and does not take the correlation with the remaining parameters into account, thus leading to lower statistical power[94]. Statistical inferences are therefore based on likelihood ratio tests as previously suggested[57,58,94]). At T1, older adults with higher LC integrity further showed steeper learning curves (likelihood ratio tests; linear slope: $\Delta\chi^2(\Delta df = 1) = 5.589$, $p = 0.018$, est = 1.717, [95% CI: 0.222, 3.212], standardized est = 0.304; quadratic slope: $\Delta\chi^2(\Delta df = 1) = 5.612$, $p = 0.018$, est = -0.360, [95% CI: -0.671, -0.049], standardized est = -0.340) in the RAVLT (see Supplementary Figure 8, left). While we did observe a reliable association between LC integrity and memory performance in older but not younger adults[13] (likelihood ratio tests: all $\Delta\chi^2(\Delta df = 1) \leq 2.691$, all $p \geq 0.101$, see Supplementary Table 8), we did not find reliable age group differences in the LC–memory association (likelihood ratio tests: all $\Delta\chi^2(\Delta df = 1) \leq 0.417$, all $p \geq 0.519$, see Supplementary Table 8).

For findings concerning the association between LC integrity and memory performance as assessed by the RAVLT at T2, please see Supplementary Table 8 and also refer to the main text and Supplementary Figure 8 (middle).

Supplementary Table 8. Overview of associations between locus coeruleus integrity and verbal learning and memory performance within and between younger and older adults for time point 1 and 2

| Covariance ($\gamma$) | TP | Age group | $\Delta\chi^2$ | $\Delta df$ | $p$ | est | CI lower | CI upper | stand. est |
|---|---|---|---|---|---|---|---|---|---|
| LC–Icept | 1 | YA | <0.001 | 1 | 0.999 | 0.003 | -3.559 | 3.566 | <0.001 |
| LC–Lin | 1 | YA | 2.691 | 1 | 0.101 | 2.418 | -0.593 | 5.429 | 0.368 |
| LC–Quad | 1 | YA | 2.425 | 1 | 0.119 | -0.497 | -1.147 | 0.152 | -0.381 |
| LC–Icept | 1 | OA | 3.993 | 1 | 0.046 | 1.256 | -0.020* | 2.532 | 0.253 |
| LC–Lin | 1 | OA | 5.589 | 1 | 0.018 | 1.717 | 0.222 | 3.212 | 0.304 |
| LC–Quad | 1 | OA | 5.612 | 1 | 0.018 | -0.360 | -0.671 | -0.049 | -0.360 |
| LC–Icept | 2 | YA | 1.181 | 1 | 0.277 | 2.014 | -1.671 | 5.698 | 0.199 |
| LC–Lin | 2 | YA | 0.796 | 1 | 0.372 | -1.062 | -3.421 | 1.297 | -0.225 |
| LC–Quad | 2 | YA | 0.802 | 1 | 0.371 | 0.200 | -0.243 | 0.643 | 0.499 |

| Covariance ($\gamma$) | TP | Age group | $\Delta\chi^2$ | $\Delta df$ | $p$ | est | CI lower | CI upper | stand. est |
|---|---|---|---|---|---|---|---|---|---|
| LC–Icept | 2 | OA | 7.939 | 1 | 0.005 | 1.737 | 0.447 | 3.027 | 0.348 |
| LC–Lin | 2 | OA | 1.033 | 1 | 0.310 | 0.637 | -0.609 | 1.882 | 0.142 |
| LC–Quad | 2 | OA | 1.426 | 1 | 0.232 | -0.167 | -0.446 | 0.112 | -0.181 |
| LC–Icept | 1 | YA vs OA | 0.417 | 1 | 0.519 | -- | | | |
| LC–Lin | 1 | YA vs OA | 0.177 | 1 | 0.674 | -- | | | |
| LC–Quad | 1 | YA vs OA | 0.146 | 1 | 0.702 | -- | | | |
| LC–Icept | 2 | YA vs OA | 0.02 | 1 | 0.889 | -- | | | |
| LC–Lin | 2 | YA vs OA | 1.588 | 1 | 0.208 | -- | | | |
| LC–Quad | 2 | YA vs OA | 1.927 | 1 | 0.165 | -- | | | |

Note: LC = Locus coeruleus; Icept = Intercept; Lin = Linear slope, Quad = Quadratic slope; TP = Time Point; YA = Younger adults (n = 66); OA = Older adults (n = 228); *df* = degrees of freedom; est = parameter estimate; CI = 95 % confidence interval; stand. est.= standardized parameter estimate; All statistical comparisons are based on likelihood ratio tests. * Please note that statistical comparisons are based on likelihood ratio tests, not confidence intervals of individual parameters (see Methods, section on cognitive data analysis)

Of note, at T2 the positive association between LC integrity and initial recall performance was also observed when analyzing younger and older adults in a common structural equation model (single group; see Supplementary Table 9).

Supplementary Table 9. Fit statistics and tests of the locus coeruleus—memory association for a single group (younger and older adults) structural equation model

| Model | Cov | $\chi^2$ | $df$ | RMSEA | CFI | $\Delta\chi^2$ | $\Delta df$ | $p$ | est | CI lower | CI upper | stand. est |
|---|---|---|---|---|---|---|---|---|---|---|---|---|
| LC–$RAVLT_{YAOA}$ | - | 50.149 | 17 | 0.082 | 0.975 | 7.538 | 1 | 0.006 | 2.047 | 0.514 | 3.579 | 0.259 |
| LC–$RAVLT_{YAOA}$ | Age | 54.821 | 20 | 0.077 | 0.976 | 7.934 | 1 | 0.005 | 1.734 | 0.466 | 3.002 | 0.219 |

*Note:* LC = Locus Coeruleus; RAVLT = Rey Auditory Verbal Learning Task; Cov = Covariate; *df* = Degrees of freedom; RMSEA = Root Mean Square Error of Approximation; CFI = Comparative Fit Index; ; est = parameter estimate; CI = 95 % confidence interval; stand. est.= standardized parameter estimate; YAOA: Younger and older adults (n = 294). Model concerns time point 2. Statistical comparisons are based on likelihood ratio tests.

Relating LC integrity to general episodic memory scores, we again observed a positive association in older but not younger adults while there were no reliable differences in this association between groups (see Supplementary Figure 9; likelihood ratio tests; older adults: $\Delta\chi^2(\Delta df = 1) = 5.108$, $p = 0.024$, $est_{older\ adults} = 0.067$, [95% CI: 0.007, 0.127], standardized $est_{older}$

$_{\text{adults}}$ = 0.331; younger adults: $\Delta\chi^2(\Delta df = 1) = 0.969$, $p = 0.325$[13], $\text{est}_{\text{younger adults}} = 0.075$, [95% CI: -0.077, 0.227], standardized $\text{est}_{\text{younger adults}} = 0.197$; age differences: $\Delta\chi^2(\Delta df = 1) = 0.01$, $p = 0.919$).

Together, these additional analyses corroborate our finding that interindividual differences in learning and memory are positively related to LC integrity in older adults (see main text) and, beyond that, suggest a stable and lasting general association.

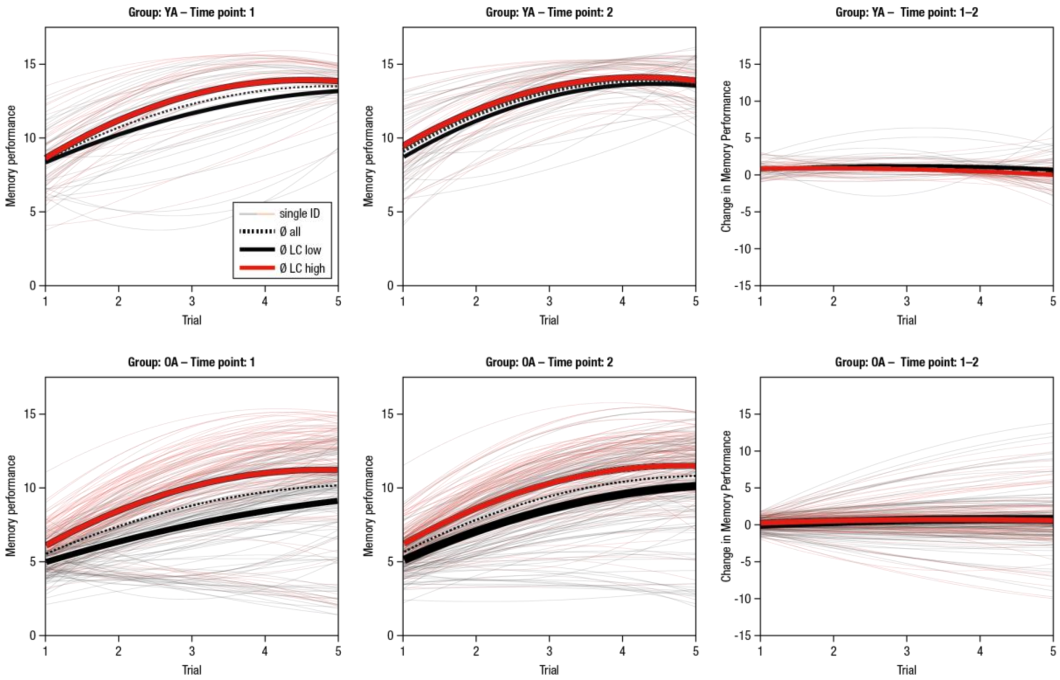

Supplementary Figure 8. Estimated learning and memory performance trajectories (RAVLT) for younger (YA; n = 66) and older (OA; n = 228) adults (upper and lower plots, respectively) for time points 1 and 2 (left and middle, respectively) and change across time points (right). For visualization of the association between locus coeruleus (LC) integrity and memory performance, single subjects (ID, thin lines) are color-coded based on LC integrity (median-split) and mean trajectories for subgroups are included (n = 33 younger adults are in each of the low and high LC groups; n = 114 older adults are in each of the low and high LC groups). Here, we used LC values automatically assessed in template space (see Methods, section on coregistration and standardization of magnetic resonance imaging data).

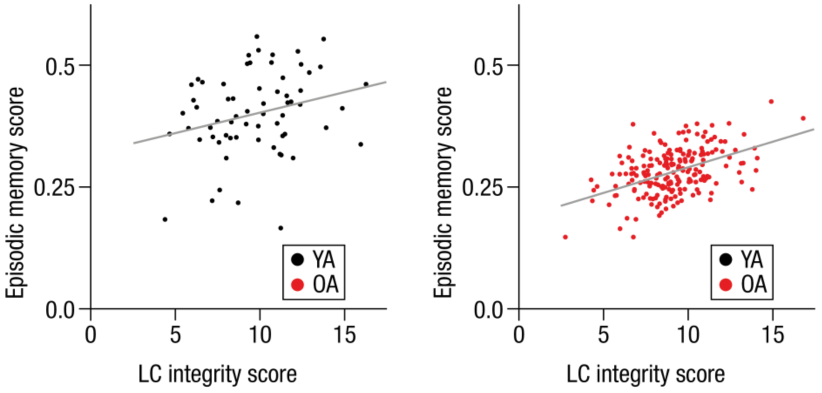

Supplementary Figure 9. Association of locus coeruleus integrity and general episodic memory performance for younger (left plot (black markers): YA; n = 66; $\Delta\chi^2(\Delta df = 1) = 0.969$, $p = 0.325$ cf.[13], $est_{younger\ adults} = 0.075$, [95% CI: -0.077, 0.227], standardized $est_{younger\ adults} = 0.197$) and older adults (right plot (red markers): OA; n = 228; $\Delta\chi^2(\Delta df = 1) = 5.108$, $p = 0.024$, $est_{older\ adults} = 0.067$, [95% CI: 0.007, 0.127], standardized $est_{older\ adults} = 0.331$) based on likelihood ratio tests (see Supplementary Figure 7).

##### 2.3.2 Associations between latent locus coeruleus integrity scores and change in memory performance

###### *2.3.2.1 Model adequacy*

The combined neural and cognitive models of each time point were then merged in a univariate latent change score model[14] (see Supplementary Figure 10). Here we only report change analyses for verbal learning and memory data (RAVLT). In doing so, we attempted to answer the question whether intraindividual (longitudinal) change in memory is related to LC scores. The model adequately represents the observed covariance matrix ($\chi^2(268) = 304.414$, RMSEA = 0.022; CFI = 0.983).

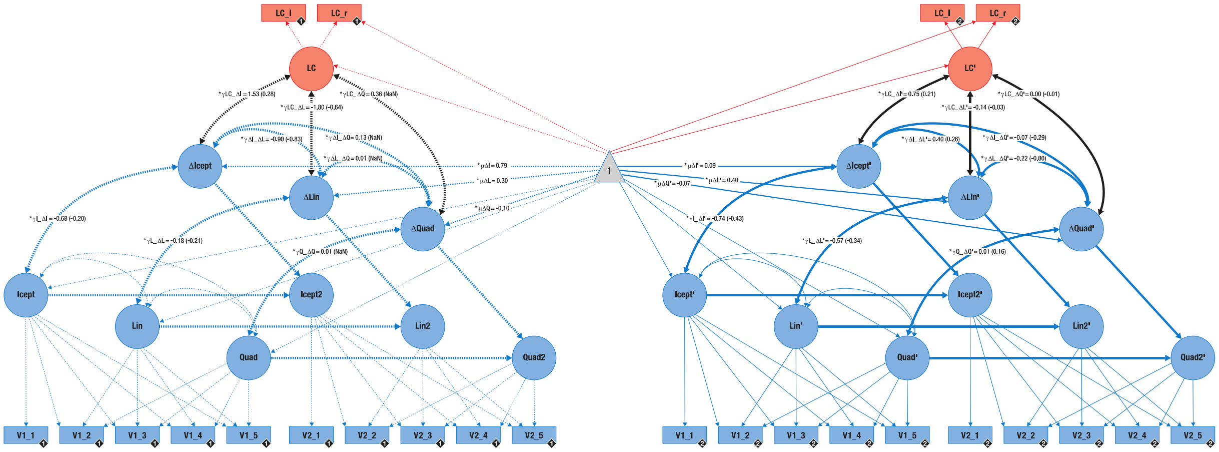

Supplementary Figure 10. Graphical depiction of the structural equation model that probes associations (black lines) between locus coeruleus integrity (LC; red) and change in memory performance (blue) in younger and older adults on a latent level. Cognitive manifest variables represent the iteratively assessed memory performance in the verbal learning and memory task (V1–5) for time point 1 (V1_*) and time point 2 (V2_*). Neural manifest variables are the LC intensity ratios of each hemisphere (left = LC_l; right = LC_r). Black diamonds on manifest variables indicate the age group (younger adults = 1, n = 66, broken lines; older adults = 2, n = 228, solid lines). (Co)Variances ($\gamma$, $\sigma$) and loadings ($\lambda$) in brackets indicate standardized estimates. Loadings that are freely estimated (*) but constrained to be equal across age groups (=) are indicated by both asterisk and equal signs (*=). Icept = Intercept; Lin/Quad= linear/quadratic slope, respectively. Rectangles and circles indicate manifest and latent variables, respectively. The constant is depicted by a triangle. Variances on manifest and latent variables are omitted to facilitate readability.

###### *2.3.2.2 Change in memory performance within younger and older adults*

In younger adults, we detected reliable average change in initial recall performance (intercept) over time (T1 → T2) as well as significant interindividual differences therein (likelihood ratio tests; mean Δ-intercept: $\Delta\chi^2(\Delta df = 1) = 7.171$, $p = 0.007$, est = 0.786, [95% CI: 0.227, 1.345]; variance Δ-intercept: $\Delta\chi^2(\Delta df = 1) = 11.986$, $p \leq 0.001$, est = 2.122, [95% CI: 0.440, 3.805]; see Supplementary Figure 8, right plots). Further, there was a trend towards change in the quadratic growth factor on the group level (likelihood ratio test: mean Δ-quadratic slope: $\Delta\chi^2(\Delta df = 1) = 3.13$, $p = 0.077$, est = -0.101, [95% CI: -0.214, 0.011]) but no reliable interindividual differences therein (likelihood ratio test: $\Delta\chi^2(\Delta df = 1) = 2.041$, $p = 0.153$, est = -

0.040 (Heywood case[14]), [95% CI: -0.088, 0.007]). We did not find any reliable mean changes in the linear slope factor (likelihood ratio test: $\Delta\chi^2(\Delta df = 1) = 1.164$, $p = 0.281$, est = 0.298, [95% CI: -0.243, 0.840]) or interindividual differences in change (likelihood ratio test: $\Delta\chi^2(\Delta df = 1) = 1.068$, $p = 0.301$, est = 0.556, [95% CI: -0.617, 1.730]).

In older adults, we observed no reliable intraindividual change (T1 → T2) in initial recall performance (likelihood ratio test: $\Delta\chi^2(\Delta df = 1) = 0.409$, $p = 0.522$, est = 0.090, [95% CI: -0.185, 0.364]) on a group level but substantial interindividual differences in the amount of change (likelihood ratio test: $\Delta\chi^2(\Delta df = 1) = 17.541$, $p < 0.001$, est = 1.304, [95% CI: 0.560, 2.048]). On average, older adults showed a trend towards intraindividual change in the quadratic slope factor (likelihood ratio test: $\Delta\chi^2(\Delta df = 1) = 3.255$, $p = 0.071$, est = -0.065, [95% CI: -0.136, 0.006]) as well as reliable interindividual differences therein (likelihood ratio test: $\Delta\chi^2(\Delta df = 1) = 5.131$, $p = 0.024$, est = 0.042, [95% CI: 0.002, 0.082]). Finally, older adults demonstrated reliable intraindividual change in linear growth and differed reliably in the amount of change they showed (likelihood ratio tests: $\Delta\chi^2(\Delta df = 1) = 5.772$, $p = 0.016$, est = 0.397, [95% CI: 0.074, 0.720] and $\Delta\chi^2(\Delta df = 1) = 31.277$, $p < 0.001$, est = 1.758, [95% CI: 0.971, 2.546], respectively).

In sum, we detected reliable intraindividual change in initial recall in younger and linear slope in older adults, indicating slightly better memory performance at the T2 in both age groups. Within the group of younger and older adults, we observed significant interindividual differences in the amount of change in initial recall and learning. However, as indicated by Supplementary Figure 8 (right), performance changes across the ~ two year interval between assessment waves tended to be negligible.

###### *2.3.2.3 Associations between latent locus coeruleus integrity scores and change in memory performance*

We did not detect reliable associations between LC integrity and change in memory performance in any of the age groups (likelihood ratio tests: all $\Delta\chi^2(\Delta df = 1) \leq 2.518$, all $p \geq 0.113$, see Supplementary Table 10). Accordingly, we do not report age group differences in these associations.

Supplementary Table 10. Overview of associations between locus coeruleus integrity and change in verbal learning and memory performance (time point 1 ) within younger and older adults

| Covariance ($\gamma$) | Age group | $\Delta\chi^2$ | $\Delta df$ | $p$ | est | CI lower | CI upper | stand. est |
|---|---|---|---|---|---|---|---|---|
| LC–ΔIcept | YA | 1.293 | 1 | 0.256 | 1.531 | -1.167 | 4.229 | 0.277 |
| LC–ΔLin | YA | 2.424 | 1 | 0.120 | -1.802 | -4.172 | 0.568 | -0.637 |
| LC–ΔQuad | YA | 2.518 | 1 | 0.113 | 0.357 | -0.105 | 0.820 | NaN* |
| LC–ΔIcept | OA | 1.921 | 1 | 0.166 | 0.747 | -0.329 | 1.823 | 0.209 |
| LC–ΔLin | OA | 0.048 | 1 | 0.826 | -0.139 | -1.381 | 1.103 | -0.034 |
| LC–ΔQuad | OA | <0.001 | 1 | 0.977 | -0.004 | -0.283 | 0.274 | -0.006 |

Note: LC = Locus coeruleus; Icept = Intercept; Lin = Linear slope, Quad = Quadratic slope; YA = Younger adults (n = 66); OA = Older adults (n = 228); *df* = degrees of freedom; est = parameter estimate; CI = 95 % confidence interval; stand. est.= standardized parameter estimate; All statistical comparisons are based on likelihood ratio tests. *Heywood case (ΔQuad), see negative variance estimate above and[14].

##### 2.3.3 Association between age differences in the spatial distribution of locus coeruleus intensity ratios and memory performance

To evaluate the functional significance of the observed topographical age differences (see main text, Figure 5), we related memory performance to LC ratios for each identified cluster. In the main text, we report analyses across all participants relying on memory performance as assessed by the RAVLT. Note that the general episodic memory model, that integrated performance over multiple memory tasks, only demonstrated weak factorial invariance (i.e., requires variant manifest intercepts across groups) and thus precluded an interpretation of age group differences in the means of latent factors[5,6]. We thus only report the association between topographical age differences and initial recall performance.

Supplementary Figure 11 shows LC ratios for younger and older adults across the rostrocaudal axis, split by performance (median split based on initial recall performance within each age group). High and low performing older adults (light and dark red lines) demonstrated differential intensity ratios in rostral LC segments that largely overlapped with the cluster identified by the age comparisons (see above for cluster results). In contrast, LC ratios of high and low performing older adults largely overlapped in caudal segments. Supplementary Figure 12 depicts the observed distributions of younger and older adults' LC intensity ratios along the rostrocaudal axis of the nucleus. Formal tests of the associations between LC sub-regions (i.e., rostral and caudal LC ratios) and memory performance (assessed by the RAVLT) across and within age groups are presented in Supplementary Table 11. Within the sub-regions, the strength of LC–RAVLT associations did not differ reliably between groups[15] (rostral: $Z = –0.999$, $p = 0.318$; caudal: $Z = –0.424$, $p = 0.672$, see Supplementary Table 11).

Recent post-mortem studies reported a gradient of (Alzheimer's) vulnerability within the LC[16,17]. In particular, the rostral and middle segments of the nucleus showed the most prominent accumulation of aggregated tau as well as cell loss across disease stages. Our findings are in line with these reports (see Supplementary Figure 13), however, since data is not in a common reference space, a direct comparison should be interpreted with caution.

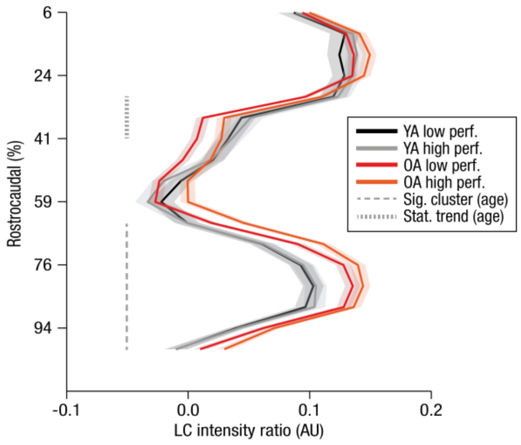

**Supplementary Figure 11**. Cluster-based permutation tests revealed spatially confined age differences in locus coeruleus (LC) intensity across the rostrocaudal axis. Age groups are split by performance (median split based on initial recall performance within each age group; n = 33 younger adults are in each of the low and high performance groups; n = 114 older adults are in each of the low and high performance groups). High and low performing older adults (light and dark red lines) demonstrated differential intensity ratios in rostral LC segments that overlapped largely with the cluster identified by the age comparisons (see broken lines). In contrast, LC ratios of high and low performing older adults largely overlapped in caudal segments. Shading represents ± 1 standard error of the mean.

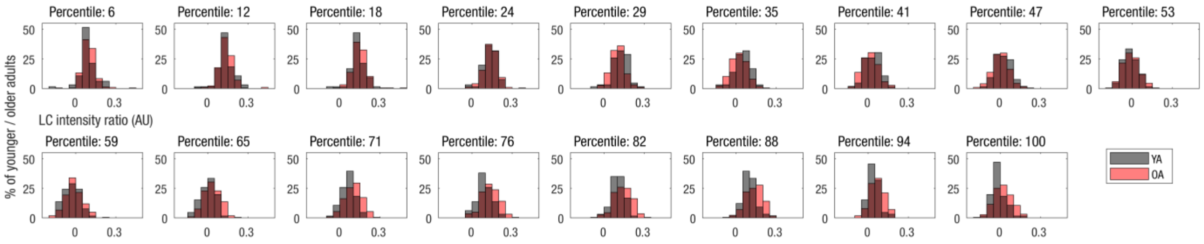

**Supplementary Figure 12**. Distributions of locus coeruleus intensity ratios along the rostrocaudal axis for younger (black; n = 66) and older (red; n = 228) adults. While older adults (OA) show higher ratios in caudal segments (> 50th percentile), there is a trend for a reversed effect in rostral segments (< 50th percentile).

**Supplementary Table 11**. Associations between rostral and caudal LC integrity and memory performance (RAVLT) within and across age groups.

| Model | Age group | $r_s$ | CI lower | CI upper | *p* |
|---|---|---|---|---|---|
| $LC_{rostral}$–$RAVLT_{intercept}$ | YA + OA | 0.207 | 0.095 | 0.314 | <0.001 |
| $LC_{rostral}$–$RAVLT_{intercept}$ | YA | 0.021 | -0.222 | 0.262 | 0.868 |
| $LC_{rostral}$–$RAVLT_{intercept}$ | OA | 0.162 | 0.033 | 0.286 | 0.014 |
| $LC_{caudal}$–$RAVLT_{intercept}$ | YA + OA | -0.08 | -0.193 | 0.035 | 0.172 |
| $LC_{caudal}$–$RAVLT_{intercept}$ | YA | 0.047 | -0.197 | 0.286 | 0.706 |
| $LC_{caudal}$–$RAVLT_{intercept}$ | OA | 0.107 | -0.023 | 0.234 | 0.108 |

*Note:* LC = Locus Coeruleus; RAVLT = Rey Auditory Verbal Learning Task; YA = Younger adults (n = 66); OA = Older adults (n = 228). CI = Confidence interval; All tests are evaluated using Spearman's correlations ($r_s$), concern time point 2 and are based on LC ratios automatically assessed in template space.

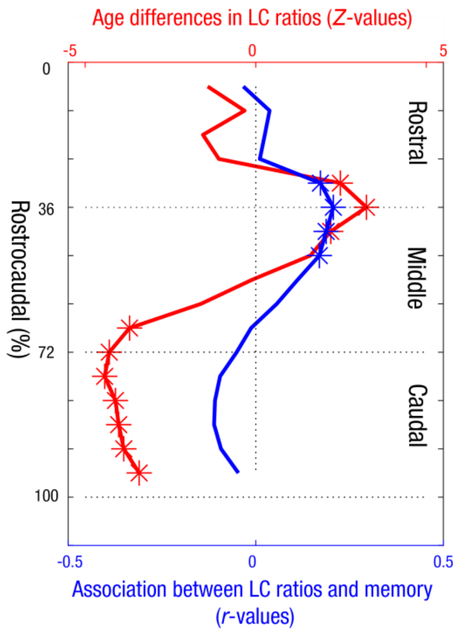

**Supplementary Figure 13.** Age differences in locus coeruleus (LC) ratios along the rostrocaudal axis (split into rostral, middle, caudal segments; red) and their association to memory performance (Rey Auditory Verbal Learning Task intercept; blue). Asterisks mark slices that showed reliable age differences (younger vs. older adults; n = 66 and n = 228, respectively); Mann-Whitney U-tests; $p < 0.05$) or associations with memory (tested across younger and older adults; n = 294); Spearman correlations; $p < 0.05$).

##### 2.4 Additional magnetic resonance imaging results

In the following, we briefly describe replications of key analyses reported above or in the main text. However, we use different LC measures here (i.e., LC values automatically extracted from native space) in order to evaluate the stability of the observed findings.

###### 2.4.1 Age differences in the spatial distribution of locus coeruleus intensity ratios

The semi-automatic procedure that we used to extract LC intensity ratios incorporates several coregistration and transformation steps (see Methods, section on coregistration and standardization of magnetic resonance imaging data) which might influence image intensity and thereby constitute a confounding factor. To exclude this possibility, we first warped the LC and reference search spaces from template space back to individual subjects' coordinates (i.e., native space) and extracted intensity values for all participants. We then calculated non-parametric cluster permutation tests[10] (see Methods, section on analysis of age differences in the spatial distribution of locus coeruleus ratios) to test for age differences in LC ratios (mean over left and right LC) across the rostrocaudal extent of the nucleus. The obtained native space results largely replicate the pattern observed in template space (see Supplementary Figure 5b, left plot): Older adults show significantly higher ratios in caudal segments (67th–83rd LC percentile; cluster permutation test: $p_{corr} = 0.002$, [95 % CI: 0.002, 0.002]), in line with the accumulation of neuromelanin across the life span[18]. In contrast, numerically, there was a reversed pattern in rostral, hippocampus-projecting segments[19]. Thus, the reported topographic analyses seem to be largely independent of transformation steps.

Studies using neuromelanin-sensitive MRI as an index for LC integrity typically reference LC signal to dorsal pontine intensity to allow for comparisons across participants[20]. However, different measures have previously been used as a basis to calculate the intensity ratio (i.e., mean, median, or max (peak) intensity)[7,8,20]. Supplementary Figure 5b (right plot) replicates the topographical analyses of age differences using mean LC and reference intensity. Compared with younger adults, older adults demonstrate elevated intensity ratios in caudal and some rostral slices (59th–82nd LC percentile; cluster permutation test: $p_{corr} = 0.008$, [95 % CI: 0.008, 0.009]; 6th–24th LC percentile; cluster permutation test: $p_{corr} = 0.017$, [95 % CI: 0.017, 0.018]) in line with age-related neuromelanin accumulation[18]. However, as observed before (see Supplementary Figure 5a), this effect is numerically reversed in the second rostral LC quartile. Thus, the mean intensity analyses largely replicated the peak intensity findings with some variations in cluster extent. Together, they point to a reduced integrity of more rostral, hippocampus-projecting LC segments[19]. Note that the mean-based LC measures produce larger (and non-negative) intensity ratios overall. Since the two previous studies that reported local age differences in LC integrity used peak intensity[7,8], we also adopt this measure for the sake of comparability.

Finally, since the intensity ratios we used do not indicate whether differences in the region of interest (LC) or reference (pons) produce the age effect, we separately assessed raw peak LC and reference intensity (see Supplementary Figure 5c). While age differences were uniformly distributed across the rostrocaudal extent (cluster permutation test: $p_{corr} = 0.002$, [95 % CI: 0.002, 0.003]) in the reference area (right plot), age differences were confined to the rostral segments (6th–53rd LC percentile; cluster permutation test: $p_{corr} = 0.019$, [95 % CI: 0.018, 0.019]; left plot) in the LC. This underscores our interpretation of reduced rostral LC integrity in late life[11].

#### 2.5 Additional combined cognitive and magnetic resonance imaging results

In the following, we briefly report replications of key analyses described above/in the main text. However, we use different LC measures here (i.e., LC values manually or automatically extracted from native space) as the basis and test their association with learning and memory (i.e., performance in the RAVLT at T2) in older adults.

##### *2.5.1.1 Model adequacy*

As described above (see Methods, section on estimation of latent locus coeruleus integrity scores), we applied SEM to estimate LC integrity scores and their relation to learning and memory. Here, however, LC integrity was calculated based on manually assessed values and then on the basis of values automatically extracted from native space (see Supplementary Figures 14 and 15, respectively). Both models demonstrate good fit (all $\chi^2(87) \leq 101.83$, all RMSEA ≤ 0.024, all CFI ≥ 0.987, see Supplementary Table 12).

Supplementary Table 12. Fit statistics for additional unified neural and cognitive models

| Model | | $\chi^2$ | *df* | RMSEA | CFI |
|---|---|---|---|---|---|
| $LC_{manual}$–RAVLT | (Time point 2) | 101.83 | 87 | 0.024 | 0.987 |
| $LC_{native}$–RAVLT | (Time point 2) | 99.208 | 87 | 0.022 | 0.989 |

*Note:* LC = Locus Coeruleus; RAVLT = Rey Auditory Verbal Learning Task; *df* = Degrees of freedom; RMSEA = Root Mean Square Error of Approximation; CFI = Comparative Fit Index.

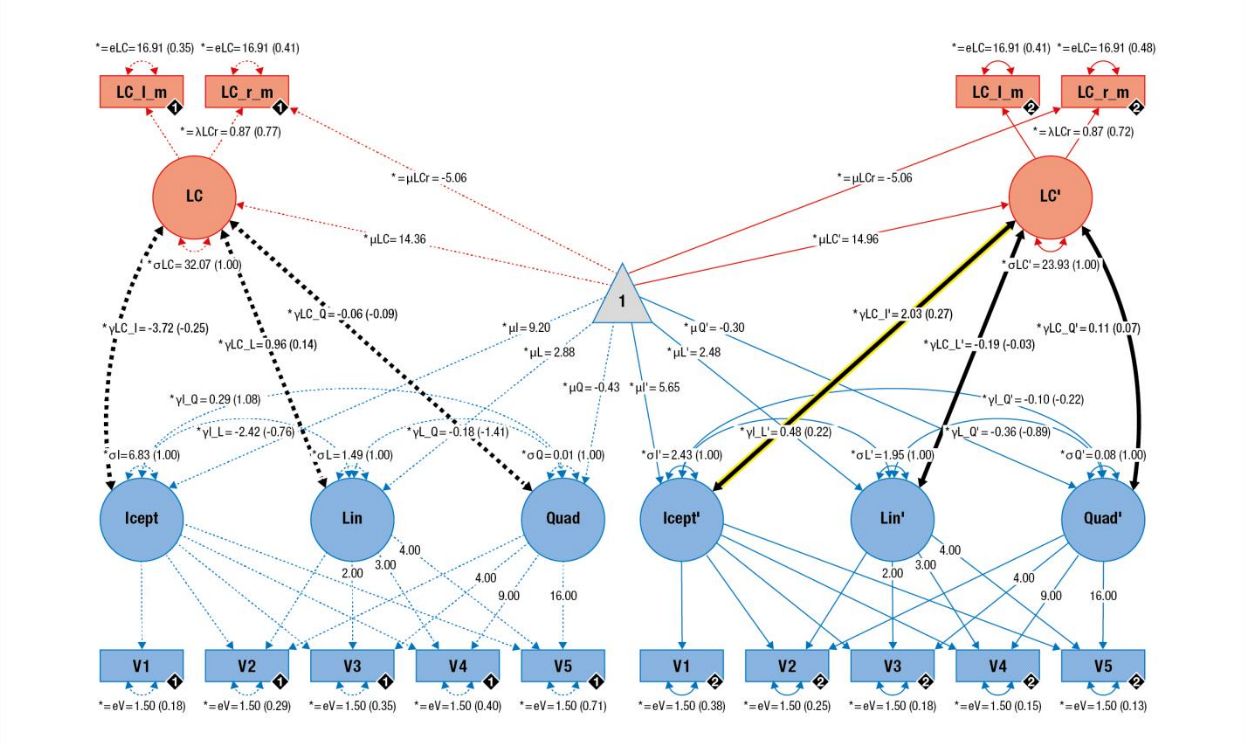

Supplementary Figure 14. Graphical depiction of the structural equation model that probes associations (thick black lines) between locus coeruleus integrity (LC; red) and learning and memory performance (blue) in younger and older adults on a latent level. Cognitive manifest variables represent the iteratively assessed memory performance in the verbal learning and memory task (V1–5). Neural manifest variables are the manually assessed LC intensity ratios of each hemisphere (left = LC_l, right = LC_r). Black diamonds on manifest variables indicate the age group (younger adults = 1, n = 66, broken lines; older adults = 2, n = 228, solid lines). (Co)Variances (*γ, σ*) and loadings (*λ*) in brackets indicate standardized estimates. Loadings that are freely estimated (*) but constrained to be equal across age groups (=) are indicated by both asterisk and equal signs (*=). Icept = Intercept; Lin/Quad= linear/quadratic slope, respectively. Rectangles and circles indicate manifest and latent variables, respectively. The constant is depicted by a triangle. The significant association between locus coeruleus integrity and memory performance (Icept) in older adults is highlighted (yellow frame).

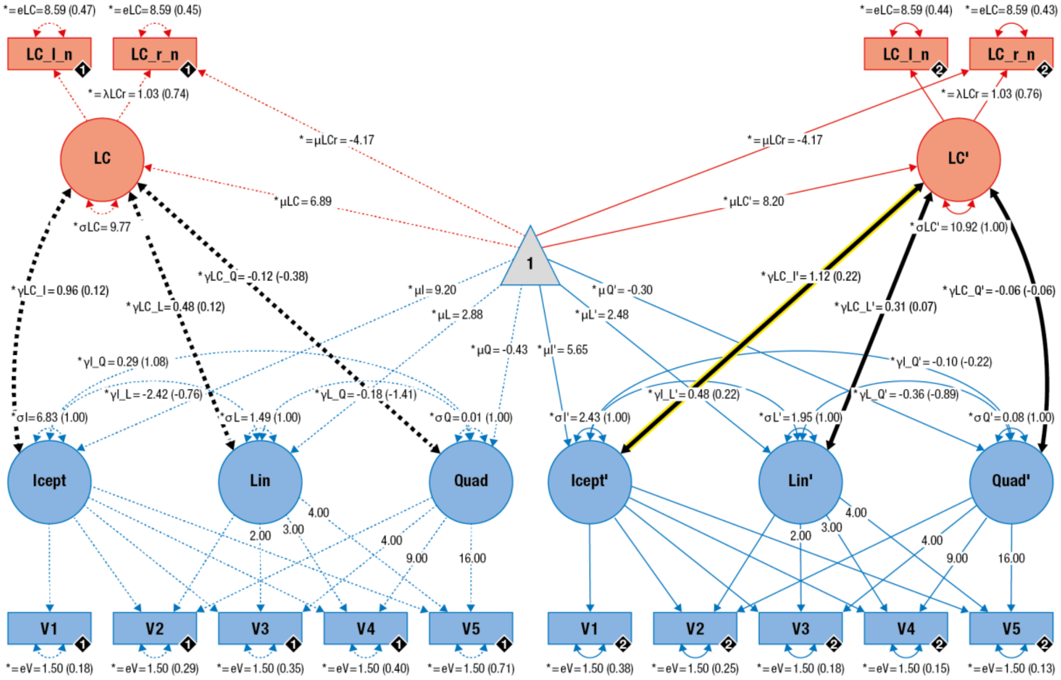

Supplementary Figure 15. Graphical depiction of the structural equation model that probes associations (thick black lines) between locus coeruleus integrity (LC; red) and learning and memory performance (blue) in younger and older adults on a latent level. Cognitive manifest variables represent the iteratively assessed memory performance in the verbal learning and memory task (V1–5). Neural manifest variables are the LC intensity ratios of each hemisphere (left = LC_l; right = LC_r), automatically assessed in native space. Black diamonds on manifest variables indicate the age group (younger adults = 1, n = 66, broken lines; older adults = 2, n = 228, solid lines). (Co)Variances ($\gamma$, $\sigma$) and loadings ($\lambda$) in brackets indicate standardized estimates. Loadings that are freely estimated (*) but constrained to be equal across age groups (=) are indicated by both asterisk and equal signs (*=). Icept = Intercept; Lin/Quad= linear/quadratic slope, respectively. Rectangles and circles indicate manifest and latent variables, respectively. The constant is depicted by a triangle. The significant association between locus coeruleus integrity and memory performance (Icept) in older adults is highlighted (yellow frame).

##### 2.5.1.2 *Latent locus coeruleus integrity scores and memory performance within older adults*

Manually segmented LC values were related to better memory performance in older adults (see Supplementary Figure 16, left). In particular, as observed before (see main results), the initial recall (intercept) factor was related to LC integrity (likelihood ratio test: $\Delta\chi^2(\Delta df = 1) = 7.541$, $p = 0.006$, est = 2.025, [95% CI: 0.539, 3.511], standardized est = 0.266).

Further, LC integrity values automatically assessed from native space showed a positive association with memory performance (see Supplementary Figure 16, right). Again, higher LC integrity was related to a higher initial recall performance (intercept; likelihood ratio test: $\Delta\chi^2(\Delta df = 1) = 5.135$, $p = 0.024$, est = 1.124, [95% CI: 0.129, 2.119], standardized est = 0.218). Taken together, these supplementary findings corroborate the positive link between LC integrity and memory in older adults across multiple analysis methods.

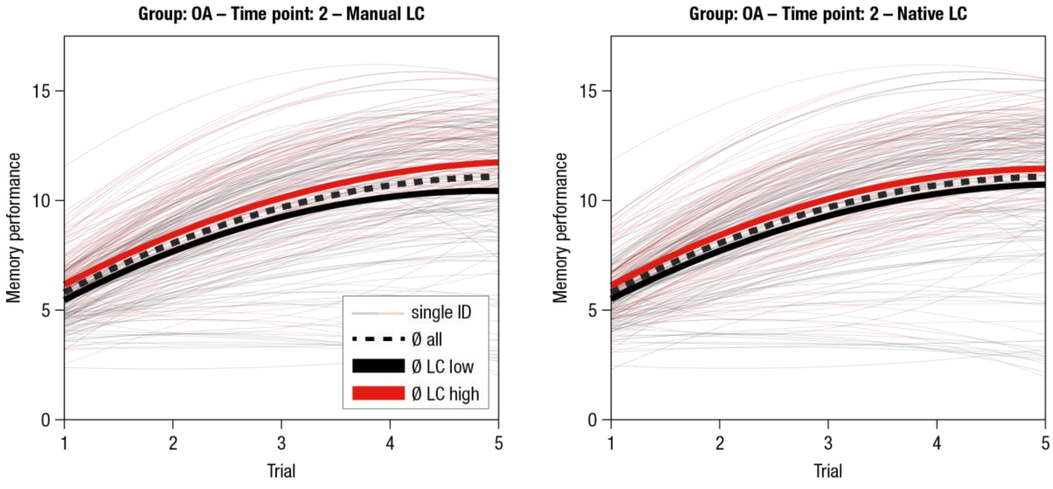

Supplementary Figure 16. Estimated learning and memory performance trajectories (RAVLT) for older adults (OA; n = 228) for time point 2. For visualization of the association between locus coeruleus (LC) integrity and memory performance, single subjects (ID; thin lines) are color-coded based on LC integrity (median split; n = 114 subjects are in the low/high LC group) and mean trajectories for subgroups are included. Left: LC integrity is calculated based on manually assessed intensity values. Right: LC integrity is estimated on the basis of automatically assessed values from native space (see Methods, section on semi-automatic locus coeruleus segmentation and intensity assessment).

#### 3 Supplementary discussion

We applied an iterative learning and memory task (RAVLT)[1] that required subjects to encode, consolidate, and retrieve verbal information and thus captures memory in its dynamic nature[2]. The non-linear performance trajectories of younger and older adults were modeled by a quadratic growth function[21] that provides a good tradeoff between parsimony and model adequacy. We used a SEM approach that structures individual differences in initial recall performance and learning. The proposed model closely approximated the observed data (i.e., excellent fit). Our analyses allowed the study of two psychologically distinct factors, namely, initial recall (i.e., performance after the first learning trial, corresponding to a standard one-trial memory assessment) and learning (i.e., changes in performance with practice). In line with previous research[3,22–24], we observed reduced initial recall performance in older adults. Age differences in the rate of learning were less consistent than in some previous reports[23], potentially due to a different statistical conceptualization of the parameter (i.e., a linear and quadratic slope factor vs. a single hyperbolic factor). However, a polynomial decomposition more accurately captured the observed performance trajectories in the present data set (see Supplementary Results 2.1.1). In their analyses of learning trajectories from eight published normative RAVLT data sets (including n = 1,222 healthy younger and older adults), Poreh[24] found that initial recall performance varied considerably by age and was influenced by demographic factors. In contrast, learning rates were highly comparable across age groups and practically independent of background factors[24]. Chronological age is, however, often considered an "empty variable" in the sense that it does not itself (but rather what it indexes) cause behavior[25]. According to the concept of brain maintenance[26,27] interindividual differences in the manifestation of age-associated brain changes and pathologies determine the amount of age-related memory decline.

In line with this notion, high levels of LC integrity may index brain maintenance that is primarily expressed in initial recall[28].

Exploiting the longitudinal nature of this data set, we evaluated performance changes over the ~ two year interval between assessment waves. However, changes in initial recall and learning factors tended to be negligible. This indicates a relative stability of memory, even in older adults. Using a comparable paradigm and population, a similar pattern emerged in the Zürich Longitudinal Study on Cognitive Aging over a delay of 1.5 years[23]. The authors mainly attributed the high stability of memory performance to retest effects resulting from a relatively short follow-up interval (see also[26], Figure 1 therein referring to[29]). Here, retest effects may even be exacerbated since identical verbal material was used at both time points. Accordingly, we mainly restricted our analyses to the neural basis of learning and memory within time points.
